# Beyond HCD: complementary fragmentation chemistries expand HLA class I immunopeptidome discovery

**DOI:** 10.64898/2026.09.24.754253

**Authors:** Terry C.C Lim Kam Sian, Fernanda Salvato, Nivedhitha Selvakumar, Tima Shamekhi, Josh Hinkle, Christopher Mullen, Gabriel A Goncalves, Ralf B Schittenhelm, Pouya Faridi

**Affiliations:** Centre for Cancer Research, Hudson Institute of Medical Research, Clayton, Australia; Department of Medicine, School of Clinical Sciences, Monash University, Clayton, Australia; Monash Proteomics and Metabolomics Platform, Clinical Proteomics Node, School of Clinical Sciences, Monash University, Clayton, Australia; Chromatography and Mass Spectrometry Department, Thermo Scientific, 355 River Oaks Pkwy, San Jose, CA 95134; Monash Proteomics and Metabolomics Platform, Monash Biomedicine Discovery Institute, Monash University, Clayton, Australia

## Abstract

Mass spectrometry based immunopeptidomics is limited by peptide identification. HLA class I ligands are non-tryptic, frequently carry low charge states and often fragment incompletely, yet nearly all workflows rely on a single activation method, higher-energy collisional dissociation (HCD). Unlike proteomics, where a protein is inferred from several peptides, in immunopeptidomics the peptide is itself the analyte and the candidate therapeutic target, so a ligand that is not identified is a target that is not found. Here we evaluate whether two alternative activation chemistries, ion trap collision-induced dissociation (CID) and infrared multiphoton dissociation (IRMPD), recover ligands that HCD does not. To our knowledge this is the first application of either method to HLA class I immunopeptidomics, made practical by the integrated infrared laser of the Orbitrap Tribrid Apex MultiOmics. Four HLA-typed cell lines were analysed in technical triplicate against the canonical human proteome and a Ribo-seq-derived cryptic ORF database. Because fragment-intensity predictors are trained almost exclusively on HCD spectra, AlphaPeptDeep was fine-tuned separately for each method; fine-tuning and Percolator rescoring increased unique peptide identifications in every cell line and method combination (median +17.1% canonical, +54.5% cryptic), and fine-tuned models transferred across cell lines with a median loss of 0.4%, placing the three methods on a common analytical footing. On that footing the three methods yielded immunopeptidomes of similar size and composition, but each recovered peptides the others missed. HCD produced the most complete fragment-ion ladders and recovered more peptides from the basic-anchor alleles A*11:01 and A*33:03, both members of the A3 supertype. CID and IRMPD recovered a larger share of alleles without a basic C-terminal anchor and, for peptides containing no lysine, arginine or histidine, produced higher search scores and more matched b/y ion current in three of four cell lines. Each method therefore contributed exclusive peptides, and at a fixed budget of three LC-MS/MS runs, one run per method yielded on average more unique peptides than triplicate runs of a single method in all four cell lines (+0.4% to +10.9%). Of 239 novel canonical predicted HLA binders in the two cancer cell lines (HCT116 and MDA-MB-231) absent from the Immune Epitope Database, 55% were detected by only one method; nine derive from cancer-associated antigens. CID and IRMPD can therefore provide sequences that may not be identified by HCD alone, or with better spectral quality for a chemically defined subset of ligands. Their value lies alongside HCD rather than in its place, and realising it will require fragment-intensity models and acquisition schemes built for these chemistries rather than adapted from HCD.

## Introduction

HLA class I molecules present intracellularly derived peptides of 8-12 amino acids at the cell surface for surveillance by CD8+ T cells, and the collective presented repertoire, the immunopeptidome, defines the antigenic landscape available for T-cell recognition [1]. Mass-spectrometry-based immunopeptidomics is the only technique that directly identifies these naturally processed and presented peptides at scale, and it has become the gold standard in basic and translational immunology [2,3]. For instance in oncology, immunopeptidomics guides neoantigen discovery, cancer-vaccine design and target selection for T-cell-engaging therapies [4,5], and large collaborative efforts have mapped tens of thousands of HLA ligands across healthy and diseased tissues [6,7]. Beyond the canonical proteome, immunopeptidomics has uncovered a large class of cryptic (non-canonical) peptides originating from supposedly non-coding regions, untranslated regions and alternative reading frames, which substantially expand the tumor-specific antigen space [8–11]. Immunopeptidomics also differs from conventional proteomics in what a single identification is worth. In proteomics a protein is usually inferred from several peptides, and any one peptide can be missed without losing the protein. In immunopeptidomics the peptide is the analyte, and it is also the candidate therapeutic target, so a ligand that is not identified is a target that is not found. Any approach that widens the set of peptides recoverable from a given sample therefore increases the chance of finding a target that can be taken forward into vaccine, T-cell receptor or T-cell engager development.

Despite this progress, mass spectrometry-based peptide identification remains the bottleneck of immunopeptidomics. A considerable fraction of HLA class I ligands are non-tryptic, frequently carry low charge states and fragment incompletely, so a large fraction of acquired tandem mass spectra go unassigned or yield too few ions to be assigned confidently [2,12]. Complementary ligand-enrichment and data-acquisition strategies deepen immunopeptidome mining [13,14], and mobile-phase additives such as dimethyl sulfoxide further enhance HLA peptide identification [15]. Currently the dominant immunopeptidomics workflows rely on a single fragmentation method, higher-energy collisional dissociation (HCD) [7,16]. Comparative studies in proteomics have long shown that no single activation method captures all peptide classes [12,17], and a few alternatives have been explored for immunopeptidomics: electron-transfer/higher-energy collision dissociation (EThcD) extends the identification depth and sequence coverage of HLA class I immunopeptidomes and can distinguish isobaric Ile/Leu residues in peptides, ultraviolet photodissociation (UVPD) complements collisional activation for de novo peptide sequencing, and integrating alternative fragmentation techniques into standard LC-MS workflows increases proteome coverage [18–21].

The available activation methods differ fundamentally in how they deposit energy. The collision-based methods, ion trap collision-induced dissociation (CID), a slow-heating method subject to a low-mass cutoff, and beam-type HCD, a higher-energy method without that cutoff, predominantly produce b/y fragment ions [12]. Electron-based methods (ETD and EThcD) generate c/z ions and preserve labile modifications [19], whereas photon-based methods deposit energy by light absorption: UVPD uses single high-energy ultraviolet photons [18], and infrared multiphoton dissociation (IRMPD) uses the sequential absorption of many infrared photons, yielding slow-heating fragmentation. Like CID, IRMPD is subject to a low-mass cutoff that depends on the precursor q value during activation, but because photoactivation is non-resonant the precursor can be held at a lower q (default q = 0.15 on the Apex, versus q ≈ 0.2–0.25 typical for peptide CID), giving a more favourable low-mass cutoff. The non-resonant nature of IRMPD activation also confers other advantageous properties versus CID, including high efficiency for singly charged precursors and sequence coverage that can exceed CID [22–25]. IRMPD has historically required specialised hardware, such as ion traps fitted with infrared lasers or Orbitrap instruments modified for photodissociation [25,26], which limited its adoption. The Thermo Scientific Orbitrap Tribrid Apex MultiOmics is the latest tribrid Orbitrap and the first to offer an integrated infrared laser, making IRMPD available alongside HCD, ion trap CID, ETD, hybrid ETD, and UVPD on a single platform [27,28]. Ion trap CID has long been used for HLA class I immunopeptidomics, and photodissociation methods have begun to be explored in this space — most notably ultraviolet photodissociation, applied to HLA-A*02:01 immunopeptides [29] — but to our knowledge no prior study has systematically compared HCD, ion trap CID and IRMPD for HLA class I immunopeptidomics, and IRMPD itself has not previously been applied to this peptide class. The chemistry of HLA ligands gives reason to expect these methods will not simply reproduce the same repertoire.

A second constraint sits downstream of acquisition. Deep learning predictors of fragment intensity and retention time, and the rescoring frameworks built on them, are now standard in immunopeptidomics and routinely recover peptide-spectrum matches that search engine scores alone leave below threshold [30–36]. These models are trained almost exclusively on HCD spectra. Any comparison of activation methods that relies on them therefore risks scoring two of the three chemistries with an HCD-shaped ruler, and would understate whatever CID and IRMPD contribute. Method-specific fine-tuning is for this reason not a refinement but a precondition for a fair comparison.

In this study we evaluate HCD, CID and IRMPD for HLA class I immunopeptidomics, acquired on the Orbitrap Tribrid Apex MultiOmics for four HLA-typed cell lines in technical triplicate. All three methods generate the same b and y ion series, but the pattern and relative intensity of those ions differ, and current identification software scores a peptide-spectrum match partly on the agreement between a predicted and an observed spectrum. Because almost all such predictors are trained on HCD data, we fine-tuned AlphaPeptDeep separately on the CID and IRMPD data and rescored with Percolator, so that each method is judged by a model of its own chemistry rather than by a proxy for HCD.

We find that the three methods return immunopeptidomes of comparable size, and that this similarity conceals a usable complementarity. HCD produced the most complete fragment-ion ladders, whereas CID and IRMPD shifted allelic representation and each recovered peptides the other methods did not, so that spreading a fixed amount of instrument time across methods increased total identifications. For peptides lacking basic residues, CID and IRMPD also produced better spectra than HCD of the same peptide. The practical consequence is that alternative fragmentation offers a route to ligands a single-method workflow does not report, and to reporting some of them with greater confidence.

## Methods

### Cell lines and HLA typing

Four HLA class I-typed human cell lines were profiled: the EBV-transformed B-lymphoblastoid cell lines BLCL064 and BLCL004, the colorectal carcinoma line HCT116 and the breast adenocarcinoma line MDA-MB-231 (MM231). HLA genotypes (**Table 1**) were determined in-house. A HeLa elastase digest was used to generate a class I-like peptide mixture for rescoring-model validation and as a transfer-learning donor; it was not HLA-eluted and was excluded from all HLA-binding analyses.

**Table 1.**
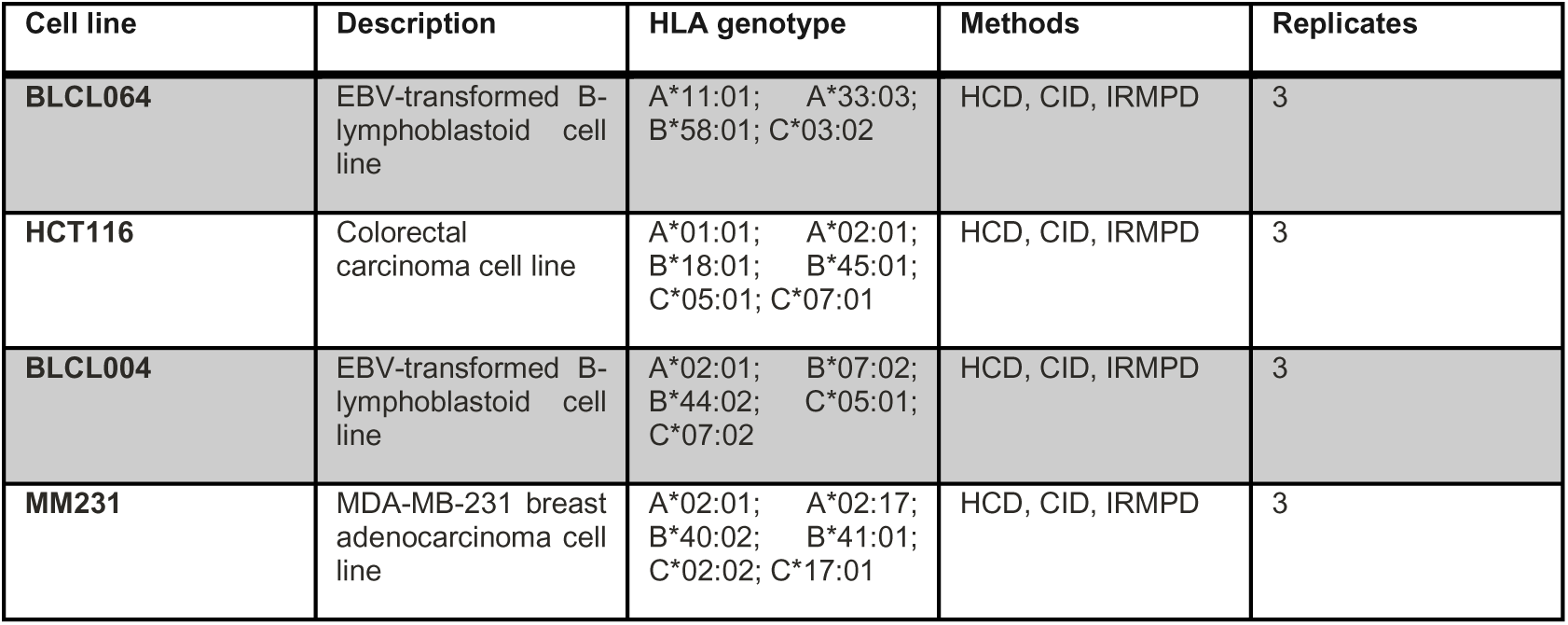
Cell lines, HLA genotypes and acquisition overview.

| Cell line | Description | HLA genotype | Methods | Replicates |
| --- | --- | --- | --- | --- |
| BLCL064 | EBV-transformed B-lymphoblastoid cell line | A*11:01; A*33:03;<br>B*58:01; C*03:02 | HCD, CID, IRMPD | 3 |
| HCT116 | Colorectal carcinoma cell line | A*01:01; A*02:01;<br>B*18:01; B*45:01;<br>C*05:01; C*07:01 | HCD, CID, IRMPD | 3 |
| BLCL004 | EBV-transformed B-lymphoblastoid cell line | A*02:01; B*07:02;<br>B*44:02; C*05:01;<br>C*07:02 | HCD, CID, IRMPD | 3 |
| MM231 | MDA-MB-231 breast adenocarcinoma cell line | A*02:01; A*02:17;<br>B*40:02; B*41:01;<br>C*02:02; C*17:01 | HCD, CID, IRMPD | 3 |

### HLA class I immunoaffinity purification

HLA class I peptide complexes were isolated by immunoaffinity purification following the SAPrIm protocol [13], a semi-automated implementation of the established W6/32 immunoaffinity workflow which was recently extended to soluble HLA immunopeptidomics (SAPrIm 2.0) [16]. Briefly, the pan-HLA class I monoclonal antibody W6/32 [37] was bound to hyper-porous magnetic protein A microbeads (MagReSyn Protein A MAX, Resyn Biosciences) and covalently crosslinked with dimethyl pimelimidate. Cell pellets were lysed in 1% (w/v) CHAPS, 50 mM Tris pH 8.0 and 150 mM NaCl supplemented with protease and phosphatase inhibitors, and HLA-peptide complexes were captured, washed and acid-eluted on a KingFisher Duo magnetic-particle processor, which handles up to 12 samples in parallel. Eluted peptides were separated from co-eluted HLA heavy chains and antibody contaminants by C18 solid-phase extraction in 96-well format (elution with 28% acetonitrile/0.1% trifluoroacetic acid), vacuum-dried, and resuspended in 2% acetonitrile/0.1% trifluoroacetic acid.

### LC-MS/MS acquisition

Peptides were analysed on a Vanquish Neo nano-LC system (Thermo Fisher Scientific; NanoCap direct-injection configuration) coupled to an Orbitrap Tribrid Apex MultiOmics mass spectrometer (Thermo Fisher Scientific) [27,28]. Separation was performed on an Ion opticks Aurora Ultimate 25cm x 75um (Ionopticks) with heating set to 50°C. Samples were loaded at 1 uL/min at 1450 bar and separated over a 72-min method with mobile phases A (water) and B (80% acetonitrile): 2% B at 0 min (0.5 uL/min), 5% B at 1 min, flow reduced to 0.2 uL/min at 1.1 min, 35% B at 61.1 min, 70% B at 63.1 min, 99% B at 67.1 min, and 99% B at 0.5 uL/min from 67.2 to 72 min.

MS1 spectra were acquired in the Orbitrap at 60,000 resolution over m/z 350-800 using vendor-default settings (AGC, injection time and RF lens per Thermo recommendation; profile mode). Data-dependent acquisition used a 2-s cycle time, peptide monoisotopic precursor selection (MIPS), charge states 1-5, and dynamic exclusion for 30 s after a single occurrence (+-10 ppm, isotopes excluded). MS/MS spectra were acquired in the Orbitrap at 60,000 resolution in centroid mode with quadrupole isolation (1.6 m/z). Fragmentation settings were: HCD, normalised collision energy (NCE) 30%; CID, NCE 36% with 5-ms activation time at q = 0.25; IRMPD, 50% normalised laser power with 3-ms activation time at q = 0.15 and 20% normalised supplemental-activation laser power (IR-TMT off). The expected peak width was set to 10 s for HCD and CID and 6 s for IRMPD. Each cell line x fragmentation method combination was acquired in technical triplicate (45 raw files in total).

### Database search

Database searching with MSFragger and downstream rescoring with AlphaPeptDeep were performed using the Biomni platform (Phylo) [38]. In brief, raw files were converted to mzML with ThermoRawFileParser (version 2.0.0.0) [39] and searched with MSFragger (version 4.4.1) [40] against two databases: first the standard human Swiss-Prot proteome (UniProt, reviewed, no isoforms; 20,431 entries, 11,418,237 residues, yielding 52,221,465 distinct 8-12mer candidate sequences under non-enzymatic search; downloaded 2025-06-19) [41]) with reversed decoy sequences, and second the cryptic database (23,230 non-canonical open reading frames derived from Ribo-seq, 845,033 residues, yielding 2,583,860 distinct 8-12mer candidate sequences; generated from the GENCODE version 45 annotation [42]) with common contaminants and reversed decoys. Searches were non-enzymatic (no cleavage specificity) with a digest mass range of 200-5,000 Da, precursor mass tolerance +-20 ppm with mass calibration and parameter optimisation enabled (calibrate_mass = 2), fragment mass tolerance 20 ppm, b and y ion series, and deisotoping and de-neutral-loss correction enabled. Variable modifications were methionine oxidation (+15.9949 Da), protein N-terminal acetylation (+42.0106 Da), pyro-Glu formation from N-terminal Gln (- 17.02655 Da) and Glu (-18.01056 Da), and Asn deamidation (+0.98402 Da), with at most two modifications per peptide; a minimum of five matched fragment ions was required per peptide-spectrum match (PSM). This modification set covers the most frequent in-vivo and sample-handling modifications of HLA class I ligands while keeping the non-enzymatic search space tractable; other modifications were profiled afterwards by open mass-offset search.

### AlphaPeptDeep rescoring and Percolator filtering

Search results were rescored with AlphaPeptDeep (peptdeep) followed by a semi-supervised linear Percolator classifier [36,43]. For each cell line x method combination, the MS/MS fragment-intensity and retention-time models were fine-tuned on all target PSMs passing 1% FDR in the pooled triplicate (peptdeep default transfer settings: 20 MS2 epochs and 40 RT epochs, each with 10 warmup epochs; fragment types b_z1, b_z2, y_z1 and y_z2 at 20-ppm MS2 tolerance; raw-specific tuning disabled); combinations with fewer than 100 training PSMs were not rescored. PSMs were re-ranked with AlphaPeptDeep’s built-in semi-supervised Percolator (a linear scikit-learn classifier; five iterations, cv_fold = 1, learning rate 0.1) and filtered to 1% PSM-level FDR by target-decoy analysis, using the identical procedure before and after rescoring so that gains are directly comparable. For the rescoring-strategy evaluation, donor models were fine-tuned analogously on (i) each other single cell line and (ii) a master pool of all five datasets (the HeLa elastase digest served as a donor only), and each donor model was applied to every recipient dataset.

### HLA-binding prediction and downstream analysis

Identified peptide sequences were queried with NetMHCpan 4.2 [44] against the corresponding cell line’s HLA genotype; peptides with eluted-ligand (EL) %Rank < 2 were considered predicted binders, and binder annotations were propagated between analyses by sequence-level lookup. Unsupervised motif analysis used GibbsCluster 2.0 [45]; clusters were assigned to HLA alleles by majority NetMHCpan best-allele vote among cluster members, and mapping purity was computed as the fraction of cluster members assigned to the cluster’s allele. Novelty was assessed against the Immune Epitope Database (IEDB) [46], and cancer-antigen candidates were mapped to a curated cancer-antigen panel. Downstream statistical analysis was performed in R (v4.4.3). Clustering was performed separately for each fragmentation method and cell line on unique 9mers (pooled triplicates, 1% FDR) with K set to the number of typed alleles per cell line and three random seeds, retaining the run with the highest final Kullback-Leibler divergence; the unionacross-methods repertoires were clustered identically.

### KRH-free peptide enumeration and fragmentation analysis

All 8-12-mer windows were enumerated from the reviewed human proteome (UniProt Swiss-Prot, 20,431 proteins; 11,418,237 residues) [41], and peptides containing lysine, arginine or histidine at any position were removed, yielding 13,629,612 unique KRH-free peptides. Binding of each KRH-free peptide to one representative allele per classical HLA supertype (A*01:01, A*02:01, A*03:01, A*24:02, B*07:02, B*27:05, B*44:02, B*58:01, B*15:01) [47,48] was predicted with MHCflurry 2.2.1 [49] (pan-allele mode; binders, elution-likelihood rank < 2). Identified ligands were assigned to the allele of best elution-likelihood rank, grouped into A2-supertype, A3-supertype and other alleles, and classified by basic-residue content as KRH-free (no K, R or H), C-terminal-basic (K or R at the C terminus) or internal-basic. For spectral-quality analysis, peptides identified by both compared methods were paired by sequence (best charge-2+ PSM per peptide), and per-peptide deltas (CID-HCD, IRMPD-HCD) of search score and summed b/y ion current were tested with two-sided Wilcoxon signed-rank tests (Benjamini-Hochberg FDR per cell line); effect sizes are median deltas with 95% bootstrap confidence intervals and rank-biserial correlations. Fragment-ion ladder completeness (fraction of theoretical b/y bonds observed) was compared between methods at fixed charge state (2+).

## Results

### Deep-learning rescoring establishes an expanded identification set for fragmentation comparison

Search-engine scores are a particular bottleneck in HLA class I immunopeptidomics. This is because HLA ligands are short, non-tryptic peptides whose fragmentation and retention time deviate from the tryptic peptides on which most scoring models are trained. Deep-learning rescoring addresses this problem directly. Specifically, AlphaPeptDeep predicts fragment intensities and retention times for each candidate peptide and re-ranks PSMs by the agreement between observed and predicted spectra. Because its models are fine-tuned on each dataset, the predictions adapt to the instrument, fragmentation method and peptide composition at hand. We expected this sample-specific training to be especially valuable here, where three fragmentation methods with systematically different fragment-ion patterns (higher-energy collisional dissociation, HCD; collision-induced dissociation, CID; and infrared multiphoton dissociation, IRMPD) are compared. Furthermore, cryptic peptides, searched against a much larger database than their canonical counterparts, stand to benefit disproportionately from spectrum-specific rescoring. As such, we first validated the fine-tuned models on the elastase-digested HeLa lysate included in the cohort as an HLA class I mimic reference, making it an ideal sample for validation.

Following rescoring, predicted MS/MS spectra closely matched their experimental counterparts. Specifically, across all FDR-passing PSMs, spectral angles between observed and predicted spectra were high for every method (median 0.93, 0.92 and 0.90 for HCD, CID and IRMPD). Notably, these were cleanly separated from decoy PSMs (per-method medians 0.16–0.24; **Figure 1a**). In addition, predicted retention times tracked observed retention times almost perfectly (Pearson r = 0.995–0.997 across methods; **Figure 1b**). This agreement is a prerequisite for trusting the identification gains reported below: only models that have genuinely learned each method’s fragmentation behaviour can be expected to re-rank PSMs beyond the search engine’s scoring. Notably, rescoring with these models increased unique-peptide identifications for the HeLa digest by 6.2–10.0% depending on the method (**Figure 1c**). Overall, these results confirm that all three fragmentation methods and the fine-tuned models performed as expected.

**Figure 1.**
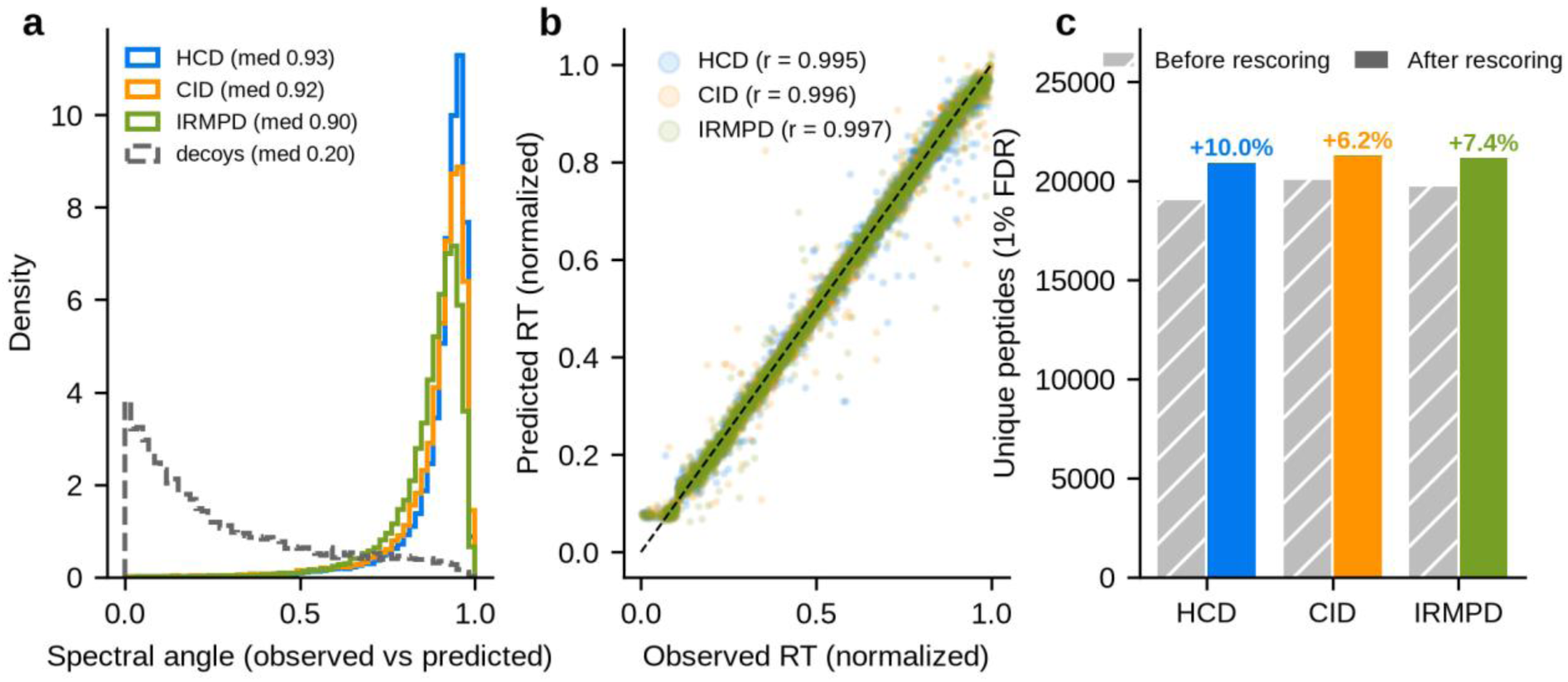
Validation of fine-tuned AlphaPeptDeep models on the HeLa elastase digest (standard database). (a) Distributions of spectral angles between observed and predicted spectra for FDR-passing target PSMs per method (solid) and for all decoy PSMs pooled across methods (dashed gray). (b) Observed versus predicted normalised retention time for FDR-passing target PSMs (4,000 randomly subsampled PSMs per method shown); r = Pearson correlation computed on all PSMs. (c) Unique peptides identified at 1% FDR before (hatched gray) and after (colored) AlphaPeptDeep rescoring of the pooled HeLa elastase triplicates per method; labels give the peptide-level gain.

Having established that the fragmentation methods and the fine-tuned models add genuine identifications, we applied the same rescoring workflow to the four HLA class I-typed cell lines (**Table 1**). Each raw file was searched against two databases in a two step search strategy on MS Fargger the standard human Swiss-Prot proteome and a cryptic (non-canonical) database generated from the GENCODE version 45 annotation [42]. All search results were then rescored with AlphaPeptDeep. In this process, models were fine-tuned on the pooled triplicates of each cell line × method combination, PSMs were re-ranked with a semi-supervised Percolator classifier, and identifications were re-filtered at 1% FDR using the same target-decoy procedure before and after rescoring. Therefore, gains are directly comparable (**Supplementary Table S1**). Unless stated otherwise, the results below refer to the standard database after rescoring at 1% FDR. The corresponding cryptic-database QC is reported in **Supplementary Figures S1–S2** and **Supplementary Tables S5–S6**.

For the standard database, rescoring increased unique-peptide identifications in all 12 cell line × method combinations. Gains ranged from +9.2% to +42.6% (median +17.1%; **Figure 2a**, **Supplementary Table S1**). Notably, gains were largest for the two cancer cell lines (MM231: up to +42.6%; HCT116: up to +34.0%). Furthermore, gains were consistent across fragmentation methods (median gain HCD +26.7%, CID +11.9%, IRMPD +17.8%).

**Figure 2.**
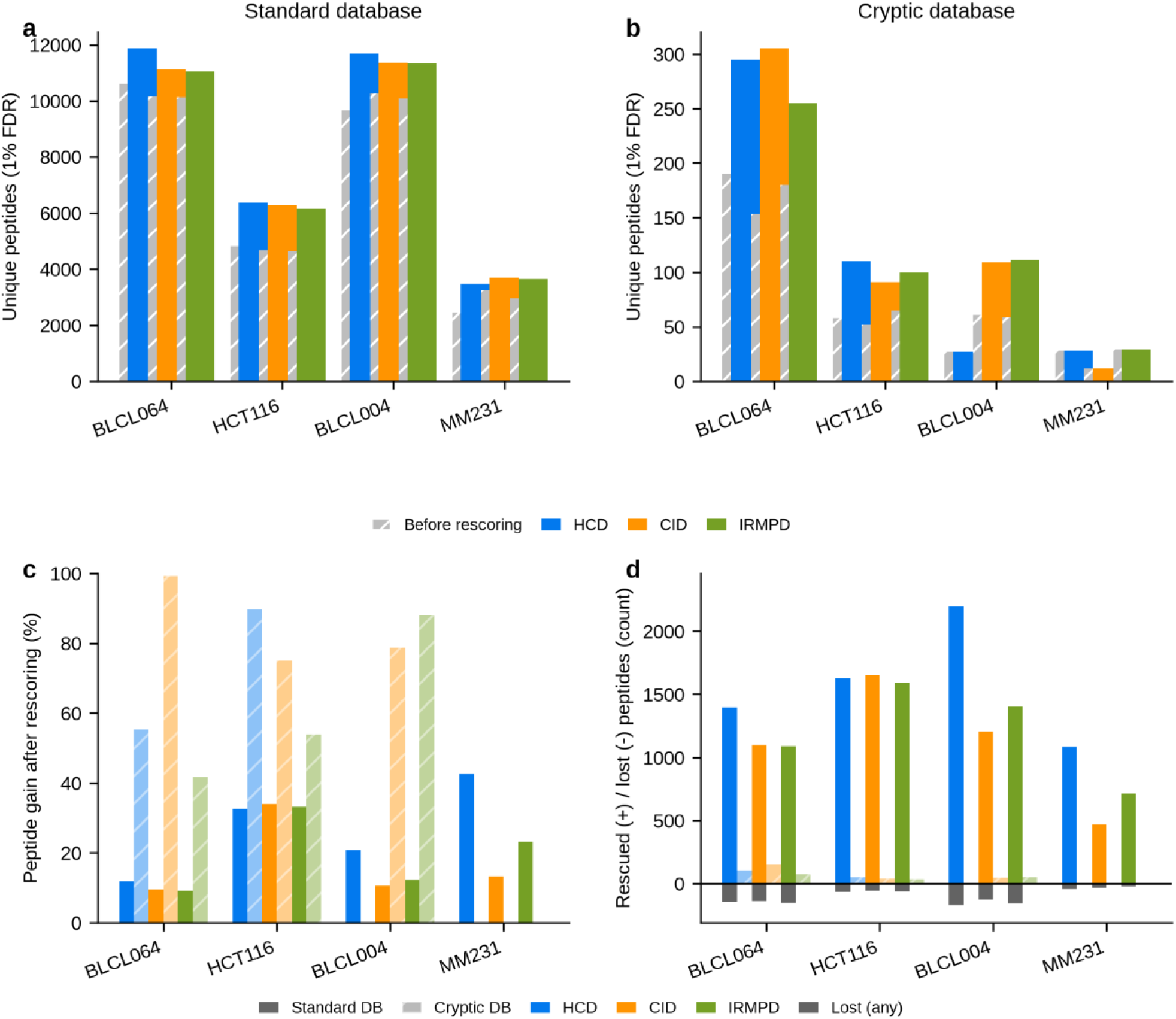
AlphaPeptDeep rescoring gains at 1% FDR. (a,b) Unique peptides before (hatched gray) and after (colored) rescoring for the standard (a) and cryptic (b) databases, per cell line and method (pooled triplicates). (c) Peptide-level gain after rescoring (% additional peptides relative to before) for both databases. (d) Rescued (newly identified, positive) and lost (no longer passing 1% FDR, negative) peptides as absolute counts; solid = standard database, hatched = cryptic database. FDR estimation used the same target-decoy procedure before and after rescoring.

Gains were substantially larger for the cryptic database (median +54.5% vs +17.1% for standard; **Figure 2b**), reaching +99.3% (BLCL064, CID). Cryptic gains were consistent across methods (median HCD +27.6%, CID +76.8%, IRMPD +47.8%). Moreover, gains were observed for every cell line with a sufficiently large cryptic repertoire. However, the four combinations with the smallest cryptic yields (the three MM231 methods and BLCL004 HCD, 12–29 peptides before rescoring) showed no net change (**Figure 2d**). The larger cryptic rescoring gain is driven primarily by the small baseline: cryptic repertoires are one to two orders of magnitude smaller than their canonical counterparts (median 89 vs 7,648 unique peptides per cell line x method), so a modest absolute number of rescued peptides produces a large percentage gain. The search space itself is not larger— the cryptic database generates 2,583,860 distinct 8-12mer candidate sequences, 20-fold fewer than the 52,221,465 from Swiss-Prot — so the higher percentage gain is a property of the baseline, not of the FDR threshold.

Peptide gains were systematic across cell lines and methods for both databases (**Figure 2c**). Importantly, re-scoring added identifications rather than merely reshuffling the list. In total, 15,521 standard-database peptides were newly identified (rescued) after rescoring, against only 1,183 lost (1.4% of the pre-rescoring repertoire; **Figure 2d**).

### Fragmentation methods produce immunopeptidomes of comparable yield, length and binding

The total number of identified peptides differed markedly between the four cell lines (**Figure 2a**, **Figure 3a, Supplementary Table S2**). The two B-lymphoblastoid lines gave the largest repertoires (BLCL064: 11,056-11,861; BLCL004: 11,327-11,685 unique peptides per method), followed by HCT116 (6,160-6,364) and MM231 (3,483-3,691). Differences between fragmentation methods within a cell line were comparatively small, with <7% between the best and worst method for every cell line. HCD gave the largest repertoires for BLCL064, BLCL004, HCT116, while CID was best for MM231. Median peptide length was 9 aa for all four cell lines (**Supplementary Table S2**).

**Figure 3.**
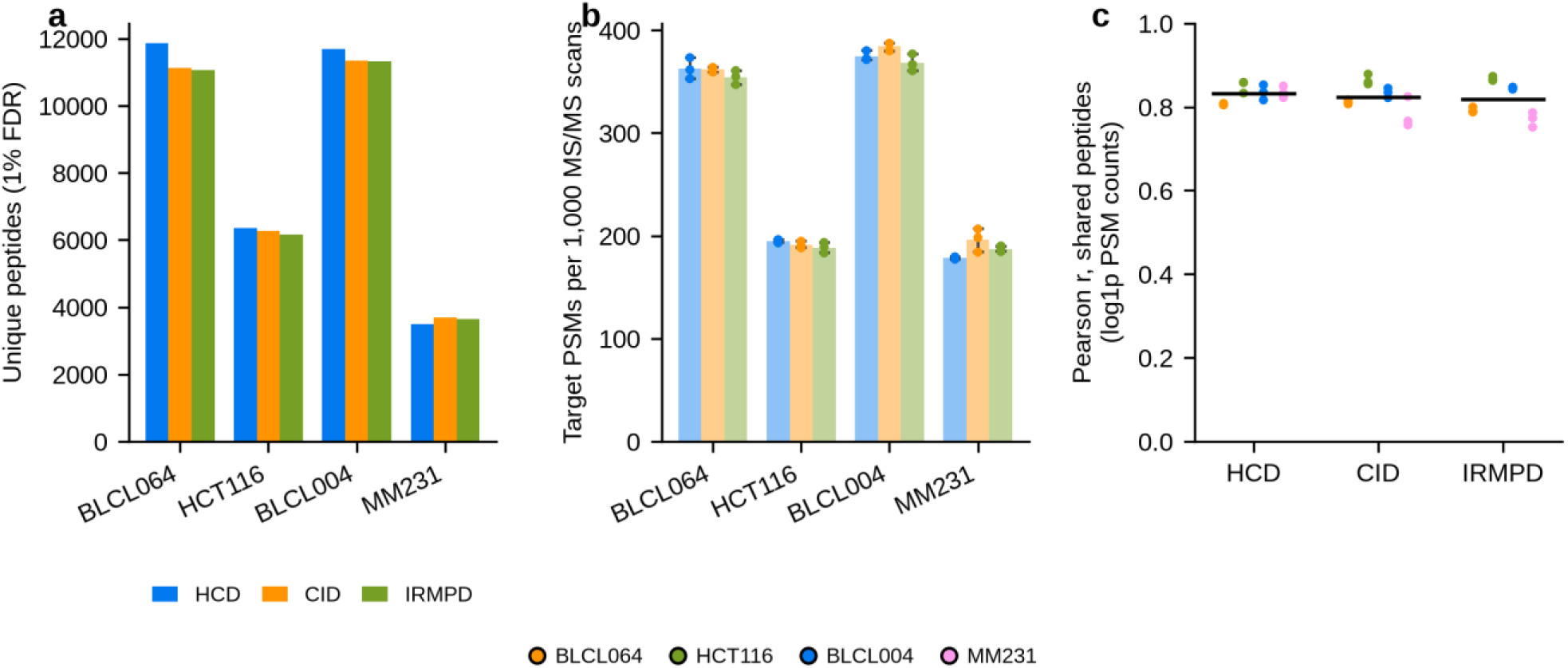
Identification quality control across the four HLA - typed cell lines and three fragmentation methods (standard database). (a) Unique peptides identified at 1% FDR after AlphaPeptDeep rescoring of pooled triplicates, per cell line and fragmentation method. (b) Identification rate: target PSMs at 1% FDR per 1,000 MS/MS scans acquired, per replicate (points) with method means (bars) and range ( error bars). (c) Pairwise Pearson correlation of log1p -transformed PSM counts for peptides detected in both replicates of a pair (quantitative agreement of the shared repertoire). Cell-line colors in (c) as in the legend below the figure. All values derive from the standard Swiss-Prot database search; cryptic-database equivalents are shown in **Supplementary Figure S1**.

Identification rates were stable across replicates and methods within each cell line and tracked overall yield, indicating that the lower yields of the cancer lines reflect lower immunopeptide content rather than acquisition artifacts. This stability is a prerequisite for the comparisons below: with identical target-decoy FDR control and reproducible acquisition across methods, differences in yield or composition can be attributed to fragmentation behavior rather than to differences in error rates or instrument performance. No fragmentation method showed a systematic identification-rate advantage (within-cell-line variation across methods <10%) (**Figure 3b**), in this regards the method choice does not trade identification efficiency for fragmentation diversity.

Technical reproducibility was assessed by quantitative correlation of the peptides detected in both replicates of a pair (Pearson r of log1p-transformed PSM counts; **Figure 3c**): mean r = 0.83, 0.82 and 0.82 for HCD, CID and IRMPD, respectively, showing that the shared repertoire is quantitatively reproducible. Cryptic-database repertoires showed the same qualitative pattern, although quantitative correlations were computed on much smaller shared peptide sets and are correspondingly noisier (**Supplementary Figure S1**).

Moreover, to confirm that the identified peptides are bona fide HLA class I ligands, we performed binding prediction of every 8 to 12mer peptide to the corresponding cell line’s HLA alleles with NetMHCpan 4.2 (EL %Rank ≤2 binder). Between 93.2% and 97.8% of peptides were predicted binders across the four typed cell lines and three methods (**Figure 4a**, **Supplementary Table S2**); cryptic-database peptides showed similarly high binder fractions (**Supplementary Figure S2**), supporting their identity as genuine HLA ligands rather than false discoveries. Assigning each binder to its best-scoring allele showed that all typed alleles contribute to the observed immunopeptidome, with HLA-A and HLA-B alleles generally presenting more peptides than HLA-C, as expected from their generally higher surface expression. Moreover, the allelic composition was nearly identical across fragmentation methods within each cell line (**Figure 4b**), showing no method-based bias. Length distributions per allele were dominated by 9mers (median 76% of an allele’s binders, range 46-91%) and were likewise consistent across methods (**Figure 4c**). To test whether fragmentation method changes the motifs recovered rather than only the number of peptides, unsupervised clustering with GibbsCluster 2.0 was performed separately on the 9mers identified by HCD, CID and IRMPD, so that the three methods can be compared directly rather than pooled (**Figure 5**). All three methods recovered allele-specific motifs. For HCT116, CID achieved one-to-one mapping of all six clusters onto the six typed alleles (mapping purity 0.49-0.98; the fraction of cluster members assigned to the cluster’s allele), while HCD and IRMPD each split A*02:01 across two clusters and absorbed one HLA-C allele. The other three cell lines showed equally specific clusters, with two recurring deviations across methods: the lowest-abundance HLA-C allele (C*03:02 in BLCL064, C*07:02 in BLCL004, C*02:02 in MM231) was frequently absorbed into a neighbouring cluster rather than recovered separately, and one allele was typically split across two sub-clusters (B*58:01 in BLCL064, A*02:01 in BLCL004, A*02:17 in MM231; **Figure 5**). Size-weighted mapping purity was 0.89-0.95 for BLCL064, HCT116 and BLCL004; MM231 was lower (0.63-0.77) because its two A2-supertype alleles A*02:01 and A*02:17 share near-identical anchor motifs and cross-map between clusters. Cluster recovery, final KLD and mapping purity overlapped across methods within each cell line, indicating that HCD, CID and IRMPD repertoires are equally allele-specific; clustering of the union-across-methods repertoires gave consistent results (**Supplementary Figure S3**).

**Figure 4.**
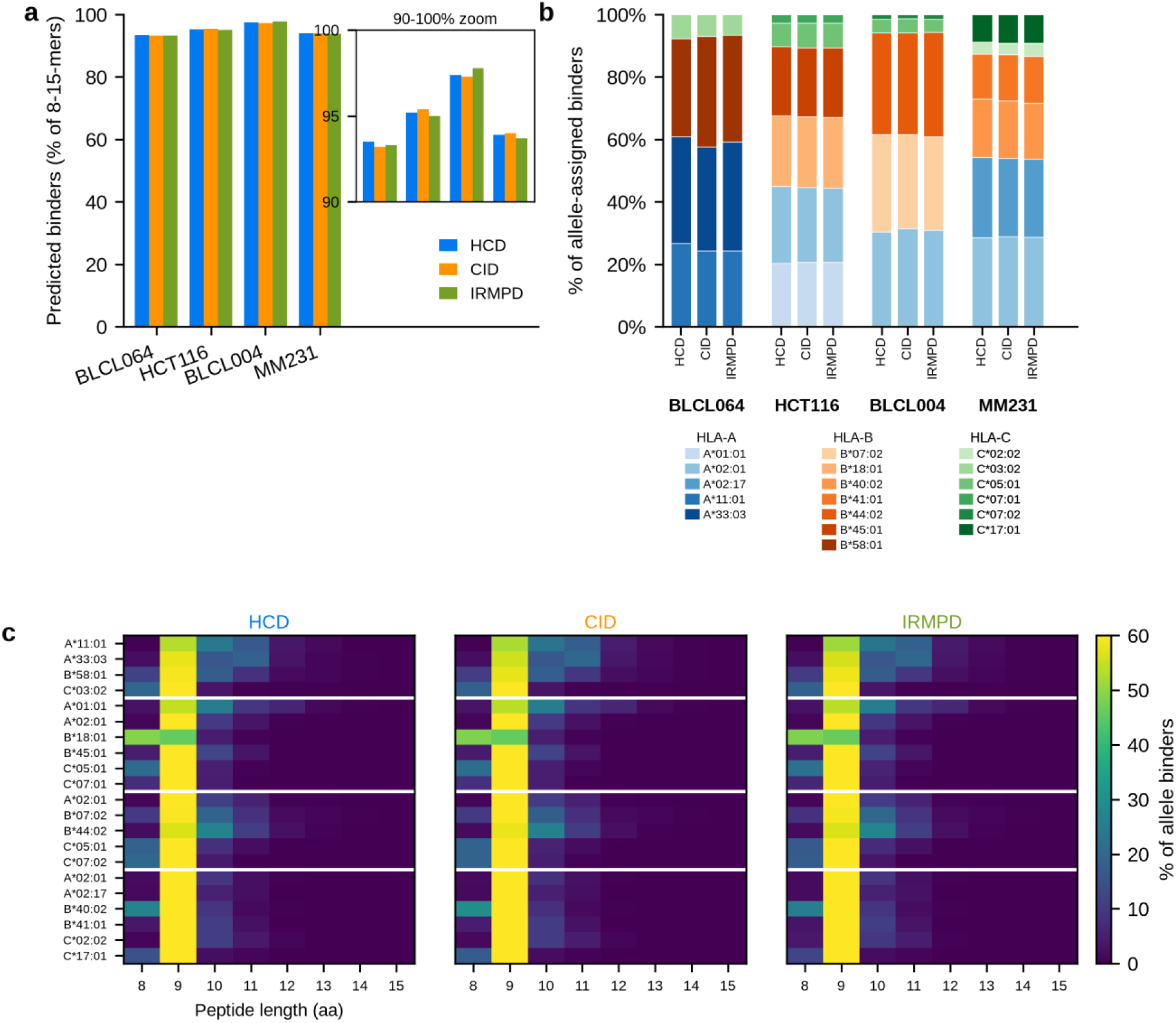
HLA-binding quality control of the identified immunopeptidomes (standard database). NetMHCpan 4.2 binding predictions for all 8-12mer peptides identified in the four HLA-typed cell lines (HeLa elastase excluded; not HLA-eluted). Binder: EL %Rank < 2. (a) Fraction of predicted peptides that are binders, per cell line and method; inset: 90-100% zoom. (b) Per-method allele contribution: binders assigned to their best-scoring allele (lowest EL %Rank), shown as the percentage of each method’s allele-assigned binders (bars sum to 100%). Colors group by locus (A: blue shades, B: orange shades, C: green shades); the legend lists all typed alleles. (c) Peptide length distribution per allele (% of the allele’s binders), one heatmap per fragmentation method; rows group ed by cell line.

**Figure 5.**
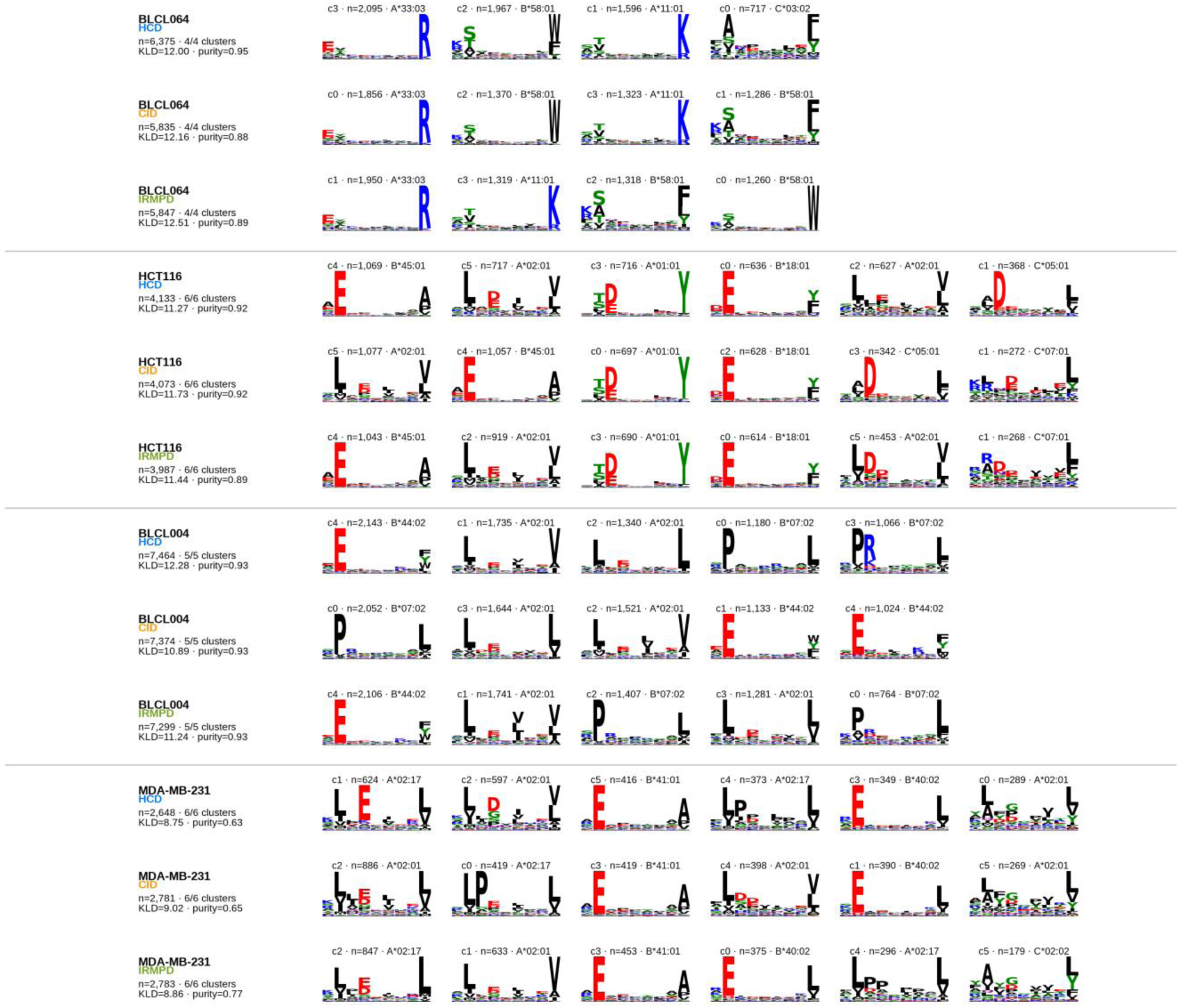
Per-method unsupervised GibbsCluster 2.0 deconvolution of the 9mer repertoires of the four HLA-typed cell lines (standard database). Unique 9mers per fragmentation method (pooled triplicates, 1% FDR) were clustered separately with K = the number of typed alleles per cell line (4-6; best of three random seeds by final KLD). Rows = cell line x fragmentation method (HCD, CID, IRMPD); columns = clusters ordered by descending size. Row annotations: number of input peptides (n), non - empty clusters/K, final KLD and size-weighted mapping purity. Information-content logos per cluster; captions give the native cluster ID, cluster size (n) and majority-mapped HLA allele. Union-across-methods clustering is shown in **Supplementary Figure S3**. HLA genotypes per cell line are listed in **Table 1**.

### HCD yields the most complete fragment-ion ladders, while CID and IRMPD shift intensity toward b-ions

To understand whether a fragmentation method gives a sequence-coverage advantage, we re-matched all theoretical b/y ions to the experimental MS/MS spectra of all pooled 1% FDR PSMs restricting the analysis to 9mer predicted binders. Peptides were assigned to one of four mutually exclusive ladder classes: both full ladders, a full b- or y-ladder only, or gap-free only (every backbone bond covered by at least one b- or y-ion without a complete series). HCD produced the highest fraction of complete ladders (36% of peptides versus 22% and 20% for CID and IRMPD, respectively) and of full b-only ladders (21%, 13% and 17%, respectively) (**Figure 6a**). Full y-only ladders were slightly more frequent in CID (28% of peptides), and IRMPD most often reached full coverage only by combining the two series (**Figure 6a; Supplementary Table S3**).

**Figure 6.**
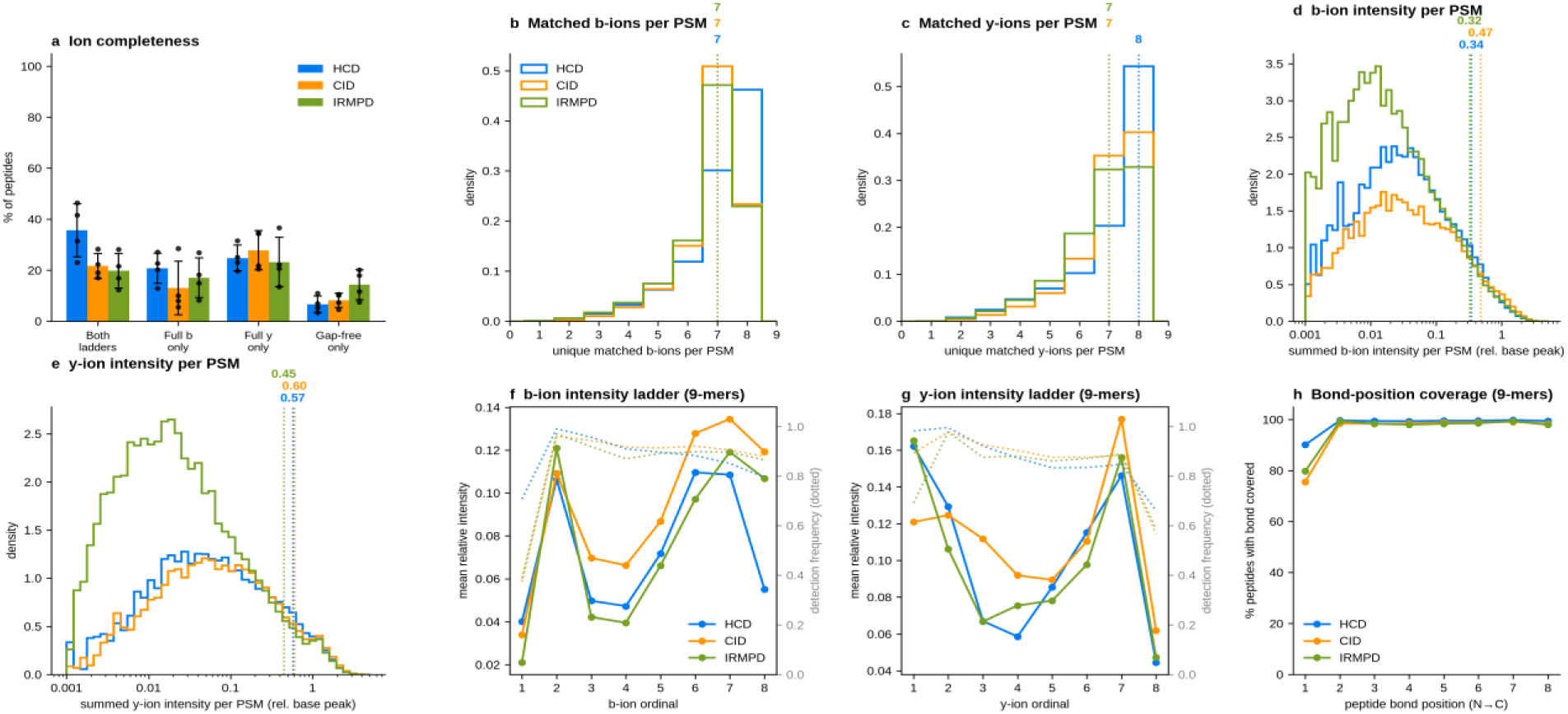
Fragment-ion coverage and intensity balance across fragmentation methods (standard database). b/y fragment ions were matched against the pooled 1% FDR PSMs of each cell line x method combination, restricting every analysis to 9mer predicted binders (best netMHCpan EL %Rank < 2). (a) Ion completeness: peptides assigned to mutually exclusive ladder c lasses - both full ladders, full b-ladder only, full y-ladder only, or gap-free without a complete series (every bond covered by at least one matched b- or y-ion); the remaining peptides (100% minus the class sum) have at least one uncovered bond; Bars = mean across the four cell lines, error bars = SD, points = individual cell lines. (b,c) Distribution of unique matched b-ions (b) and y-ions (c) per PSM: density-normalised step histograms (integer bins), PSMs pooled across the four cell lines; color = method; vertical dotted lines mark the pooled-distribution medians (labeled). (d,e) Same for summed matched-ion intensity per PSM for b-ions (d) and y-ions (e) (log10 scale, relative to the base peak). (f,g) Mean relative intensity (solid) and detection frequency (dotted) per fragment ordinal for b-ions (f) and y-ions (g) of the 9mer binders, averaged over the four cell lines. (h) Bond-position coverage: % of 9mer-binder peptides with at least one matched ion flanking each backbone bond position (N - to C-terminus), averaged over the four cell lines. Cryptic-database equivalents are in **Supplementary Figure S4** ; per-cell-line distributions in **Supplementary Figure S5**; per-cell ordinal ladders in **Supplementary Figure S6**.

The two ion series contributed matched ions in equal numbers but not equal intensity (**Figure 6b,c**). The per-PSM distributions of unique matched ions per series overlapped almost perfectly between the b- and y-series for all three methods (**Figure 6b,c**), at roughly 7-8 unique ions per series per PSM (median b-ions 7 for all three methods; median y-ions 8 for HCD versus 7 for CID and IRMPD; **Supplementary Figure S5**). In contrast, the ion intensity, was carried predominantly by y-ions for every method, the y-series intensity distributions sitting systematically above the b-series ones, even at the individual cell line level **(Figure 6d,e** and **Supplementary Figure S5**; median b-share 0.30-0.50 across cell lines and methods), and this balance shifted toward b-ions for CID and IRMPD relative to HCD (BLCL064 0.39 versus 0.30; MM231 0.49 versus 0.41). The CID shift arose from gained b-ion intensity (pooled median 0.47 versus 0.34 for HCD) at essentially unchanged y-ion intensity (0.60 versus 0.57), whereas the IRMPD shift reflected lost y-ion intensity (0.45 versus 0.57 for HCD) at unchanged b-ion intensity (0.32 versus 0.34). HCT116 again showed no difference (b-share 0.41-0.42 for all three methods) (**Figure 6d,e, Supplementary Figure S5**).

Deconvoluting the intensity profiles of the 9-mer peptides resolved the origin of these shifts (**Figure 6f,g**): CID and IRMPD spectra have roughly two-fold more intense b8-ions (0.12 and 0.11 vs 0.06 mean relative intensity for HCD), and CID additionally shows stronger b3-b4 ions (0.07 vs 0.05), whereas HCD concentrates intensity in y1 (0.16 vs 0.12 for CID). This is consistent with the known propensity of beam-type CID for C-terminal basic-residue y1 formation and of resonant CID for sequential b-ladder formation [50,51]. Bond-position coverage maps of 9mers confirmed that the CID/IRMPD completeness deficit is concentrated at the N-terminal-most bond (position 1 covered for 90% of HCD peptides versus 75% for CID), with all other positions covered for >98% of peptides by every method (**Figure 6h**). The same trends held for the cryptic database (**Supplementary Figure S4**), and the corresponding per-cell profiles were highly consistent (**Supplementary Figure S6**). We note that cross-method intensity comparisons are influenced by instrument settings (collision energy, injection time) and should be read as within-method b-versus-y balance rather than absolute sensitivity.

### Fragmentation-method-dependent amino-acid and allelic preferences, and the compositional footprint of rescoring

Despite similar global sequence composition, fragmentation methods differed in their relative coverage of individual HLA-associated repertoires. Position-resolved amino-acid frequencies of pooled 9mers differed between CID and HCD by at most 1.1 percentage points (K@P9) and between IRMPD and HCD by at most 1.3 percentage points (K@P9) (**Figure 7a,b**). Every method independently recovered the expected HLA anchor structure, with enriched P2 and C-terminal residues (**Supplementary Figure S7**). Dipeptide-level enrichment was likewise highly correlated between methods, with modest but reproducible differences concentrated in cysteine- and tryptophan-containing dipeptides (CID vs HCD: CS enriched +1.03, CL depleted -1.07 log2 units, mean across cell lines; **Supplementary Figure S8**), plausibly reflecting method-dependent detectability of carbamidomethyl-cysteine-containing peptides. CID and IRMPD repertoires were also mildly enriched for P2 glutamate and leucine (E@P2 +0.7/+0.8 pp, L@P2 +0.7/+0.8 pp for CID/IRMPD vs HCD; **Figure 7a,b**). Thus, at the level of bulk and position-resolved sequence composition the three methods sample essentially the same immunopeptidome.

**Figure 7.**
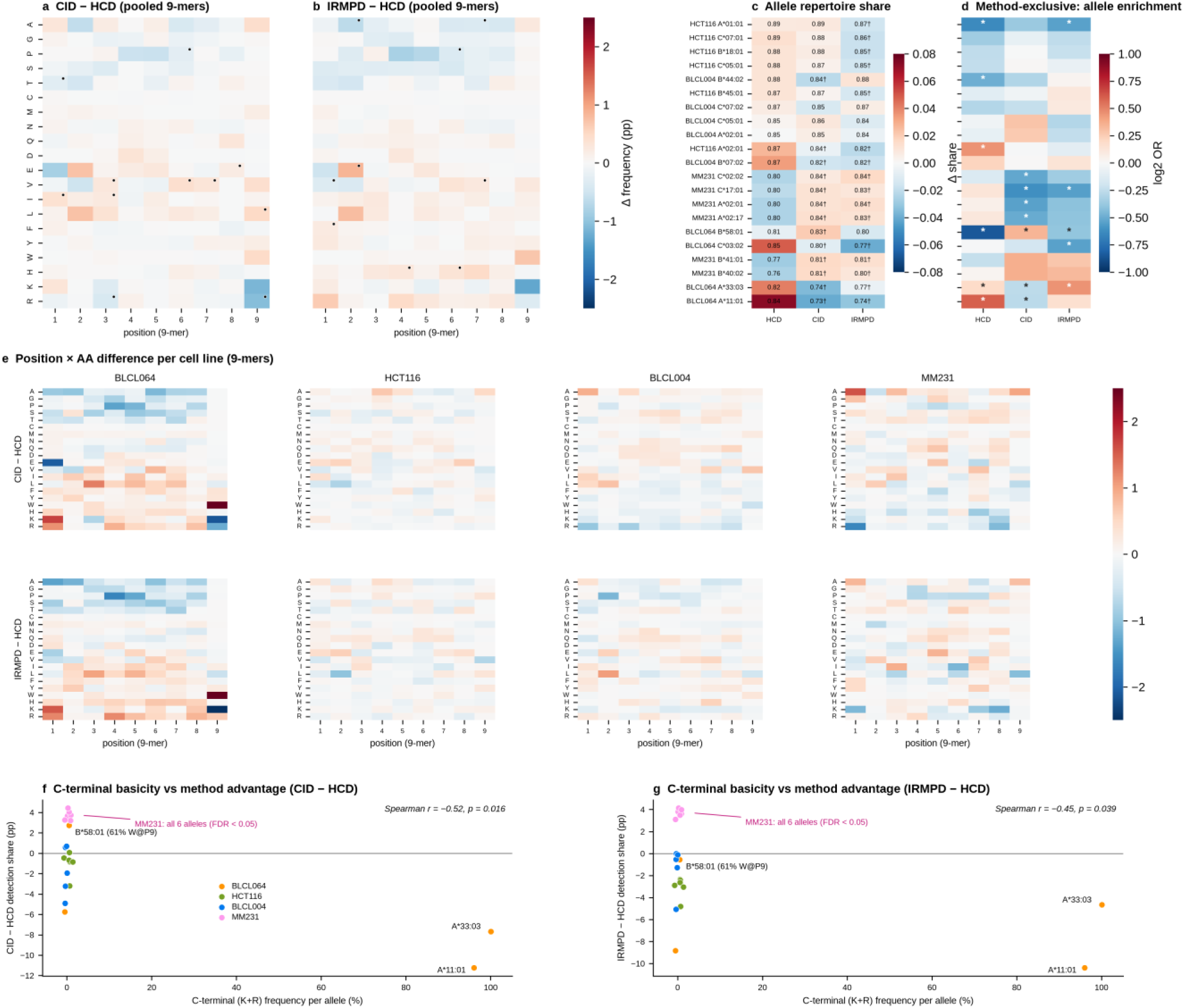
Position-specific amino-acid and allelic preferences of the three fragmentation methods (standard database). (a, b) Position x amino-acid frequency difference (percentage points) for pooled 9mers (union across the four HLA-typed cell lines), CID minus HCD (a) and IRMPD minus HCD (b); red = higher frequency in the first-named method; dots mark cells with a consistent direction of change across all four cell lines (pooled |delta| >= 0.2 pp). (c) Per-allele identification share: fraction of each allele’s predicted-binder repertoire (union across methods) detected by each method. Color = deviation from the per-allele mean share; printed numbers = raw share; † = significant difference from HCD (two-proportion z-test, Benjamini-Hochberg FDR per cell line, FDR < 0.05; full statistics in Supplementary Table S4). (d) Allelic enrichment of method-exclusive peptides (detected by only one method): log2 odds ratio per allele, Fisher exact test with Benjamini-Hochberg FDR per cell line; * FDR < 0.05; rows are in the same order as panel c. (e) Position x amino-acid frequency difference (as in a,b) broken down per cell line. (f) C-terminal basicity versus method advantage for CID: each point is one allele (colored by cell line; x-axis jittered for visibility) plotting the allele’s combined K+R frequency at P9 against its CID minus HCD detection-share difference; Spearman r = -0.52, p = 0.016 (n = 21 alleles). (g) As in (f), for IRMPD minus HCD; Spearman r = -0.45, p = 0.039. HeLa elastase is not HLA-typed and is excluded. Per-method position-specific enrichment against the proteome background is in Supplementary Figure S7, dipeptide-level enrichment in Supplementary Figure S8, per-allele position-specific differences in Supplementary Figure S 9, and method-exclusive sequence motifs in **Supplementary Figure S10** ; cryptic-database equivalents in **Supplementary Figure S11**.

Differences emerge at the cell-line and allele level. The one substantial position-specific signal, a pooled C-terminal lysine difference (K@P9), was itself a composition effect rather than a motif change. This difference was concentrated entirely in BLCL064 (CID-HCD −2.1, IRMPD-HCD −2.8 percentage points), with the other three lines flat (**Figure 7e**). It tracked allele representation, not any change in presented sequence. Specifically, HCD detected 84% of A*11:01 binders versus 73% (CID) and 74% (IRMPD) (**Figure 7c**), and A*11:01 is a lysine-anchored allele (88% K@P9). Therefore, HCD’s higher K@P9 frequency in the pooled repertoire simply reflects its over-sampling of A*11:01. Within A*11:01 itself, the K@P9 frequency was essentially identical across methods (88–89%). More broadly, per-allele position-specific profiles were flat for every allele with sufficient counts (maximum within-allele difference 4.6 pp; **Supplementary Figure S9**). As such, fragmentation method shifts which alleles are sampled but does not alter the motif any allele presents.

Systematic per-allele testing confirmed that these sampling shifts are significant and reciprocal (**Figure 7c,f**; **Supplementary Table S4**). Of the 21 alleles tested, 13 (CID) and 16 (IRMPD) differed significantly from HCD in detection share. Notably, HCD’s advantage was concentrated on the two A3-supertype alleles: A*11:01 (−11.2 and −10.4 pp for CID and IRMPD, FDR < 1e-30) and A*33:03 (94% R@P9; −7.7 and −4.7 pp). Conversely, CID and IRMPD each recovered a significantly larger share of all six MM231 alleles (+3.2 to +4.4 pp, all FDR < 0.05). CID additionally recovered more of BLCL064 B*58:01 (+2.7 pp, FDR < 0.001). B*58:01 is the dominant tryptophan-anchored allele (61% W@P9, constant across methods). Therefore, the BLCL064 W@P9 excess in CID and IRMPD repertoires (+3.1 and +2.5 pp, FDR < 0.05; **Figure 7e**) is the mirror image of the A*11:01 lysine effect. Across alleles, C-terminal basicity (combined K+R frequency at P9) correlated with the CID-HCD share difference (Spearman r = −0.52, p = 0.016; **Figure 7f**), and similarly for IRMPD (r = −0.45, p = 0.039; **Figure 7g**). In contrast, no P2 anchor residue showed such a significant association (E@P2, L@P2, P@P2, A@P2 all p > 0.1). However, the correlation attenuated when the two basic-anchor alleles were excluded (r = −0.31, p = 0.19), and A*02:01 flipped direction across cell lines (HCD-favoured in HCT116, CID/IRMPD-favoured in MM231). As such, the basicity axis should be read as a strong but partly context-dependent trend. Within these limits, HCD’s weak spots are precisely where CID and IRMPD recover more. Therefore, at the allele level, the methods are complementary.

Method-exclusive peptides sharpened the allelic picture and confirmed the complementarity. In BLCL064, HCD-exclusive peptides were enriched for A*11:01 (log2 OR +0.59, FDR < 0.05) and depleted for B*58:01 (-0.85), while CID-exclusive peptides showed the reciprocal B*58:01 enrichment (+0.34, FDR < 0.01; IRMPD-exclusive peptides were B*58:01-depleted, -0.39) (**Figure 7d**, per-cell-line values in **Supplementary Figure S12**); A*11:01 and B*58:01 thus form a complementary pair in which each method recovers peptides the others miss. In MM231 all four significantly enriched alleles among CID-exclusive peptides were depleted (log2 OR -0.61 to -0.50), indicating an excess of non-binders among CID-only identifications in that line. Consistent with the allelic mechanism, method-exclusive peptides carried distinctive motifs: HCD-exclusive 9mers were strongly enriched for C-terminal lysine and arginine (K@P9 +11.7, R@P9 +11.5 percentage points) and depleted for P2 glutamate and leucine (E@P2-12.1, L@P2 -7.6), whereas CID- and IRMPD-exclusive peptides showed no comparably strong motif (all |delta| <= 4.8 pp; **Supplementary Figure S10**) - the CID/IRMPD advantage is broad allele-repertoire coverage rather than a specific sequence chemistry. Allelic and motif preferences therefore exist but are cell-line-specific rather than universal, and pooling methods remains the safest strategy for unbiased allele coverage. The same qualitative patterns held for the cryptic database (**Supplementary Figure S11**).

Finally, we asked whether AlphaPeptDeep rescoring mitigates or introduces compositional bias by comparing rescued, lost and common peptide sets (**Figure 8**, **Supplementary Table S5**). Rescoring strongly mitigated binding-specificity bias: peptides lost after rescoring were predominantly non-binders (only 34% NetMHCpan binders versus 96% for the common set), while rescued peptides remained overwhelmingly binder-like (90%). Rescoring did, however, introduce a subtle but reproducible sequence-composition footprint: rescued peptides were depleted of aromatic and sulfur-containing residues (W-0.86, Y -0.59, F -0.43, M -0.47 log2 OR versus common) and enriched for small aliphatic residues (A +0.24, L +0.31, I +0.22), consistently across methods and cell lines (Figure 8b), the expected behaviour for a rescoring model that adds the most evidence where search-engine scores are weakest. Method-specific shifts were also observed: HCD preferentially rescued hydrophobic, doubly charged peptides (median GRAVY +0.24 vs -0.34 for common; 13% charge ≥ 3+ vs 21%), whereas CID rescued higher-charge peptides (34% vs 16%) (medians across the four cell lines, as in **Supplementary Table S5**; **Supplementary Figure S13)**. Lost peptides were on average longer (median 10 aa vs 9 aa) and less likely to carry a basic C-terminus (**Figure 8a)**. Net compositional drift is therefore small relative to the gain in binder purity, but should be considered when comparing rescued-only peptide sets across methods.

**Figure 8.**
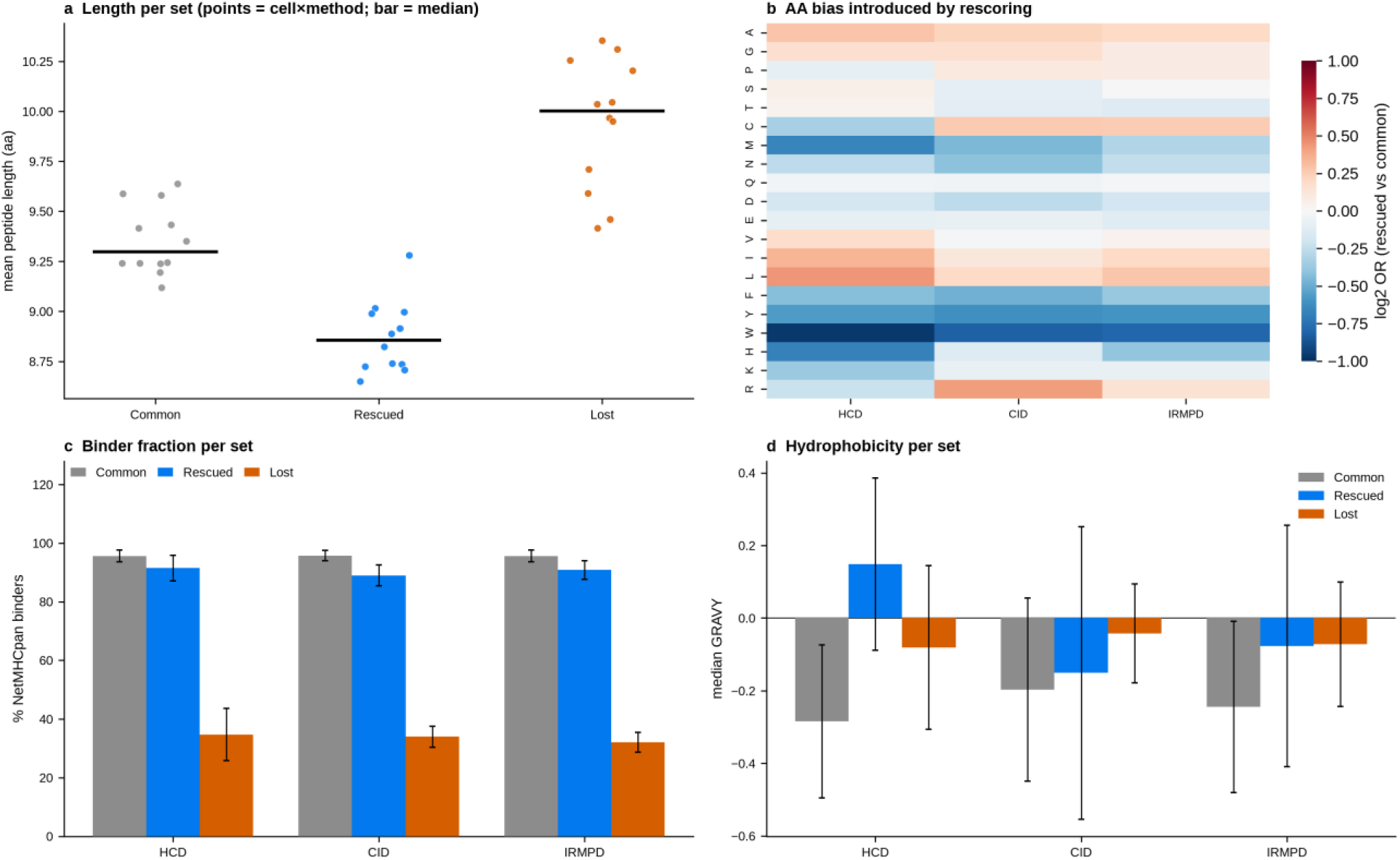
Compositional footprint of AlphaPeptDeep rescoring (standard database). Peptides were partitioned per cell line x method into common (passing 1% FDR before and after rescoring), rescued (passing only after) and lost (passing only before). (a) Peptide length per set (points = cell line x method means; bar = median). (b) Single-amino-acid log2 odds ratio of rescued versus common peptides, mean across the four cell lines, per method. (c) NetMHCpan 4.2 binder fraction (EL %Rank < 2, best allele) per set and method, mean +- SD across the four HLA-typed cell lines; binder calls include supplementary predictions for peptides absent from the primary prediction tables. (d) Median GRAVY hydrophobicity per set and method, mean +- SD across the four cell lines.

Finally, we profiled post-translational modifications across methods with an open mass-offset search of the pooled 1% FDR PSMs (standard database, four HLA-typed cell lines; **Supplementary Figure S14**). Phosphorylated (S/T/Y) peptides were recovered most efficiently by HCD and CID and were depleted under IRMPD (median 0.48% and 0.30% versus 0.14% of PSMs; q = 0.056 and 0.056, borderline after FDR control). Lysine acetylation showed the opposite preference and was significantly enriched under IRMPD (0.011% vs 0.000% for HCD, q = 0.043), with lysine/arginine methylation following the same trend without reaching significance (q = 0.138); diGly ubiquitination-remnant peptides were recovered best by HCD (0.034% vs 0.000% for CID, q = 0.043). Met oxidation (∼7% of PSMs) and deamidation (∼0.7% of PSMs) were unchanged across methods (all q > 0.1). Beyond modifications, the overall physicochemical profiles of the identified repertoires - length, hydrophobicity, charge, isoelectric point and sequence-composition metrics - were dominated by cell-line identity rather than fragmentation method (**Supplementary Figure S15**).

### Methodological breadth can outperform replicate depth at the same instrument-time budget

Having established that the three fragmentation methods differ in fragment-ion coverage, allelic sampling and sequence motifs, we quantified the overlap between their identifications in order to determine how instrument time could best be allocated. The methods sampled substantially non-redundant immunopeptidomes, with 57.2-71.1% of peptides recovered by all three methods (**Figure 9a**). Since a substantial proportion of peptides was not shared by all three methods, we next analysed how identification yield changes with each added replicate and/or method (**Figure 9b**). Notably, identification gains diminished with each addition: in BLCL064, for instance, a second HCD replicate added 13.7% more unique peptides, but a third added only 6.4% (9,799 -> 11,143 -> 11,861), and the same pattern was observed for CID and IRMPD in all four cell lines (**Figure 9b**). Therefore, at a fixed budget of three LC-MS/MS runs, splitting the budget across methods (one replicate each of HCD, CID and IRMPD) yielded more unique peptides on average than three replicates of a single method, with substantial gains in three of the four cell lines: +10.9% in BLCL064, +4.5% in BLCL004 and +4.3% in MM231 (mean cross-method yield versus mean single-method triplicate; Figure 9b). The gain was negligible in HCT116 (+0.4%), whose 71% all-method core leaves little for breadth to add. Cryptic-database equivalents showed the same pattern, with larger gains in three lines (+18.8% BLCL064, +32.0% BLCL004, +22.5% MM231) and a smaller gain in HCT116 (+3.3%; **Supplementary Figure S16**)

**Figure 9.**
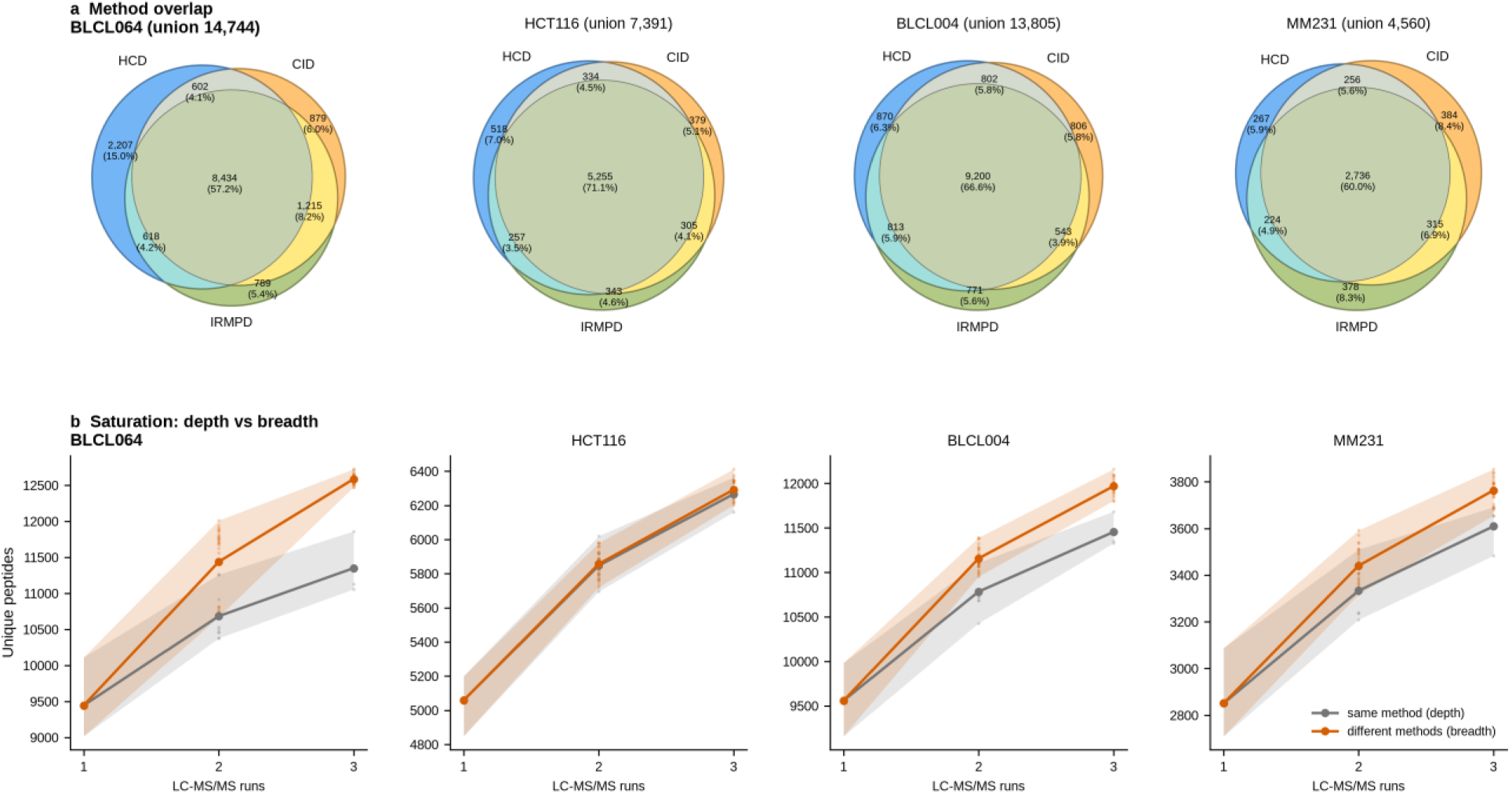
Non-redundant sampling and saturation analysis of the three fragmentation methods (standard database). (a) Three-way overlap of the unique peptides identified by HCD, CID and IRMPD (pooled triplicates, 1% FDR after rescoring) per cell line; numbers give peptides per partition (percent of the union in parentheses) and the union size. (b) Saturation of unique-peptide yield with instrument time: grey = same-method accumulation (mean over replicate subsets), orange = cross-method accumulation (mean over method x replicate allocations); shaded bands = range, points = individual allocations. Cryptic-database equivalents are in **Supplementary Figure S16**.

### Novel HLA ligands and cancer-antigen candidates

To assess whether the multi-method immunopeptidomes contain previously unreported HLA ligands, we matched every identified peptide against the HLA class I human peptides curated in IEDB. In the two cancer cell lines, 3% to 5% of identified peptides were absent from IEDB, giving 155 novel predicted binders in HCT116 and 84 in MM231 (EL %Rank < 2; **Figure 10a**). More than half of these 239 novel binders — 131 (55%) — were detected by exactly one fragmentation method (37 HCD-only, 46 CID-only, 48 IRMPD-only; **Figure 10b**). Methodological breadth is therefore directly linked to novel ligand discovery: a single-method workflow would not have reported these peptides at all. Nor are the three contributions interchangeable: HCD-exclusive novel binders were enriched for peptides assigned to A*02:01 and A*02:17, the two A2-supertype alleles that dominate both cancer lines, whereas CID-exclusive and IRMPD-exclusive novel binders were distributed across the remaining alleles without a basic C-terminal anchor (**Supplementary Tables S9 and S10**). HCD therefore contributes novel ligands from a narrow and predictable chemical niche, while CID and IRMPD contribute novel ligands spread across the remainder of the repertoire; as a result, the two together cover more of the presented landscape than either does alone.

**Figure 10.**
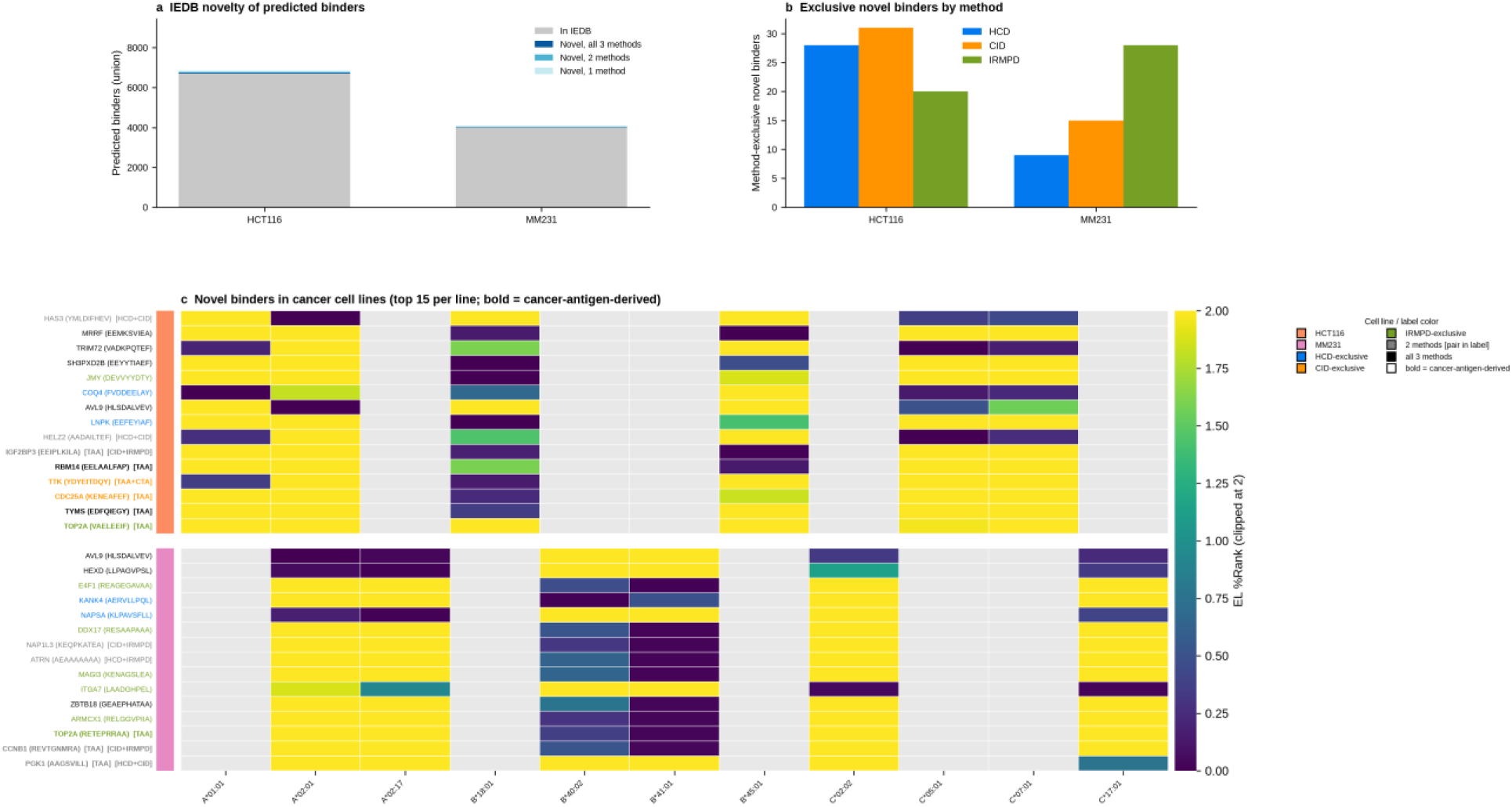
Novel HLA ligands and cancer-antigen candidates (standard database). Peptides were matched against 1,033,961 unique human peptides curated in IEDB; binders are NetMHCpan 4.2 EL %Rank < 2. (a) Predicted binders (union across methods) in the two cancer cell lines (HCT116, MM231), partitioned into in-IEDB (grey) and novel (blue shades, by number of detecting methods); the equivalent panel for the two non-malignant reference lines (BLCL064, BLCL004) is shown in Supplementary Figure S18. (b) Method-exclusive novel binders per cancer cell line, colored by detecting method (reference lines in Supplementary Figure S18). (c) Novel binders in the two cancer cell lines: all nine cancer-antigen-derived novel binders (bold, labeled with the cancer-antigen gene and category tag [TAA]/[CTA]/[TAA+CTA]) plus the top-ranked remaining novel binders, 15 per cell line (ranked by best EL %Rank): EL %Rank per typed allele (clipped at 2; dark = lowest EL %Rank; light grey = allele not present in that cell line). Row labels give the source gene and peptide sequence, colored by detecting-method set (for peptides detected by two methods the pair is named in brackets in the label); the left strip marks the cell line. Full novel-binder lists are provided as Supplementary Tables S9 and S10; cryptic equivalents in **Supplementary Figure S17.**

Turning to clinical relevance, intersecting the novel binders of the two cancer lines with a curated cancer-antigen panel identified nine cancer-antigen-derived novel binders, six in HCT116 and three in MM231 (8 TAA, 1 TAA+CTA; **Figure 10c, Supplementary Table S6**). Four of the nine were detected by a single fragmentation method, and two more were found by CID and IRMPD but not HCD; as such, only three of the nine would have been identified by an HCD-only workflow. The strongest candidates included IGF2BP3 EEIPLKILA [TAA] (EL %Rank 0.008, CID and IRMPD), TOP2A RETEPRRAA [TAA] (EL %Rank 0.029, IRMPD-only) and CCNB1 REVTGNMRA [TAA] (EL %Rank 0.034, CID and IRMPD); the top cancer-testis candidate was TTK YDYEITDQY [TAA+CTA] (EL %Rank 0.144, CID-only) (**Figure 10c**).

The cryptic immunopeptidome, by contrast, was far less covered by IEDB: 42.9% to 81.0% of cryptic peptides per cell line and method were absent from it. In the two cancer lines, this yielded 74 novel predicted binders (HCT116 55, MM231 19), of which 29 (39%) were detected by a single fragmentation method (**Supplementary Figure S17**). Fragmentation dependence is therefore stronger for cryptic than for canonical ligands: two-fifths of the novel cryptic binders rest on one activation method alone, so a cryptic antigen discovery workflow restricted to HCD would return a correspondingly smaller candidate list. As expected for non-canonical reading frames, none mapped to the curated cancer-antigen panel. The full novel-binder lists, including cancer-antigen annotations, are provided as **Supplementary Table S7 (**standard) and **Supplementary Table S8** (cryptic).

### A master rescoring model per fragmentation method matches per-dataset performance across cell lines

We next asked whether a fine-tuned model can be transferred across cell lines, which would remove per-dataset fine-tuning altogether. Across the 48 single-donor transfers (five donors, the four HLA-typed lines plus the non-tryptic HeLa elastase digest, × four HLA-typed recipients, excluding self-pairs), the donor model recovered a median of 99.6% of the peptides obtained with the recipient’s own model (5th percentile 91.3%; **Figure 11a**). HCD transferred essentially without loss (0.994-1.001 for all donor-recipient pairs). The clear exception was HCT116 in CID and IRMPD: as a recipient, foreign donors recovered only 90.3-92.3% (CID) and 92.7-94.7% (IRMPD) of its own-model repertoire, and as a donor HCT116 was itself the weakest model (mean 96.7% for CID, 93.6% for IRMPD), including the global minimum of 88.9% (HCT116 -> MM231, IRMPD). Independent repeat fine-tuning on the same data reproduced the own-model result to within 0.2% (Jaccard >= 0.9918), so these deficits are genuine model differences rather than training noise. Notably, the HeLa elastase model, fine-tuned on a non-tryptic digest of a different cell type, transferred within 0.7% to every non-HCT116 recipient, indicating that the fine-tuned models predominantly learn fragmentation behavior rather than sample-specific biology.

**Figure 11.**
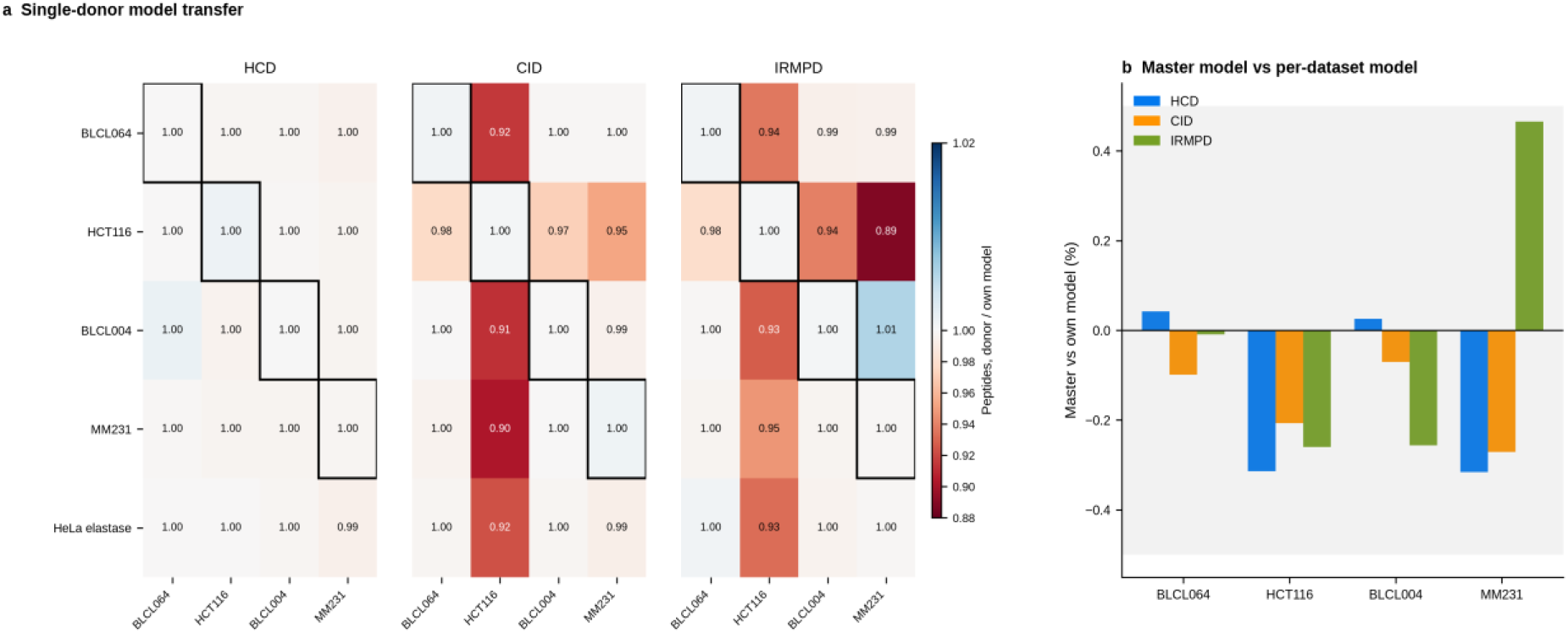
Rescoring strategy evaluation: cross - cell-line model transfer and a master rescoring model (standard database). (a) Single-donor transfer: unique peptides after rescoring with a model fine-tuned on the donor cell line (rows) relative to the recipient’s own model (columns), per method; boxed diagonal cells are self-transfer controls (independent repeat fine-tuning on the same data). HeLa elastase (non-tryptic digest) is included as a donor only. (b) Master-model rescoring relative to the per- dataset own model for the 12 cell line x method combinations; grey band = ±0.5%.

A single master model per fragmentation method, fine-tuned on the pooled PSMs of all five cell lines, performed within +-0.5% of the own-data model for all 12 cell line × method combinations (ratio 0.9968-1.0047; **Figure 11b, Supplementary Table S9**), with essentially unchanged NetMHCpan binder fractions (-0.01 to +0.32 percentage points) and high peptide-level overlap (Jaccard >= 0.983). The master model was also the only non-self donor to solve HCT116 CID and IRMPD (99.8% and 99.7% of own-model peptides), because the recipient is part of its training pool.

### KRH-free peptides couple A2-supertype anchor chemistry to a CID/IRMPD spectral-quality advantage

Two earlier findings pointed to basic-residue content as a key determinant of fragmentation. First, the allelic complementarity in **Figure 7**: HCD favored basic-anchor alleles, whereas CID and IRMPD favored hydrophobic ones. Second, the y-dominance of C-terminal-basic peptides in all three methods (**Figure 6**). We therefore asked how large the class of peptides entirely lacking basic residues would be in the HLA-presented repertoire, and whether such peptides show a systematic fragmentation-method advantage. To answer this, we enumerated every possible 8–12 amino-acid peptide from the reviewed human proteome (UniProt Swiss-Prot, 20,431 proteins; 11.42 million residues) and retained only peptides containing no lysine, arginine or histidine at any position, yielding 13,629,612 unique KRH-free peptides. We then predicted their binding across the nine classical HLA supertypes, retaining 2,620,243 predicted binders (19.2%; rank < 2). Notably, the yield tracked anchor chemistry: the A2 supertype, whose C-terminal anchor accepts hydrophobic residues, alone accounted for 638,499 predicted binders (4.7% of all KRH-free peptides), whereas the A3 supertype, whose members (including A*11:01 and A*33:03) require a C-terminal lysine or arginine, retained only 32,851 (0.24%) by construction (Figure 12a). The eluted ligand data confirmed this bias: 23–37% of A2-assigned ligands were KRH-free in the three A2-positive lines (HCT116 33.1%, BLCL004 36.8%, MM231 22.6%), versus 0.2% of A3-assigned ligands in BLCL064 (**Figure 12b**; **Supplementary Table S10**).

**Figure 12.**
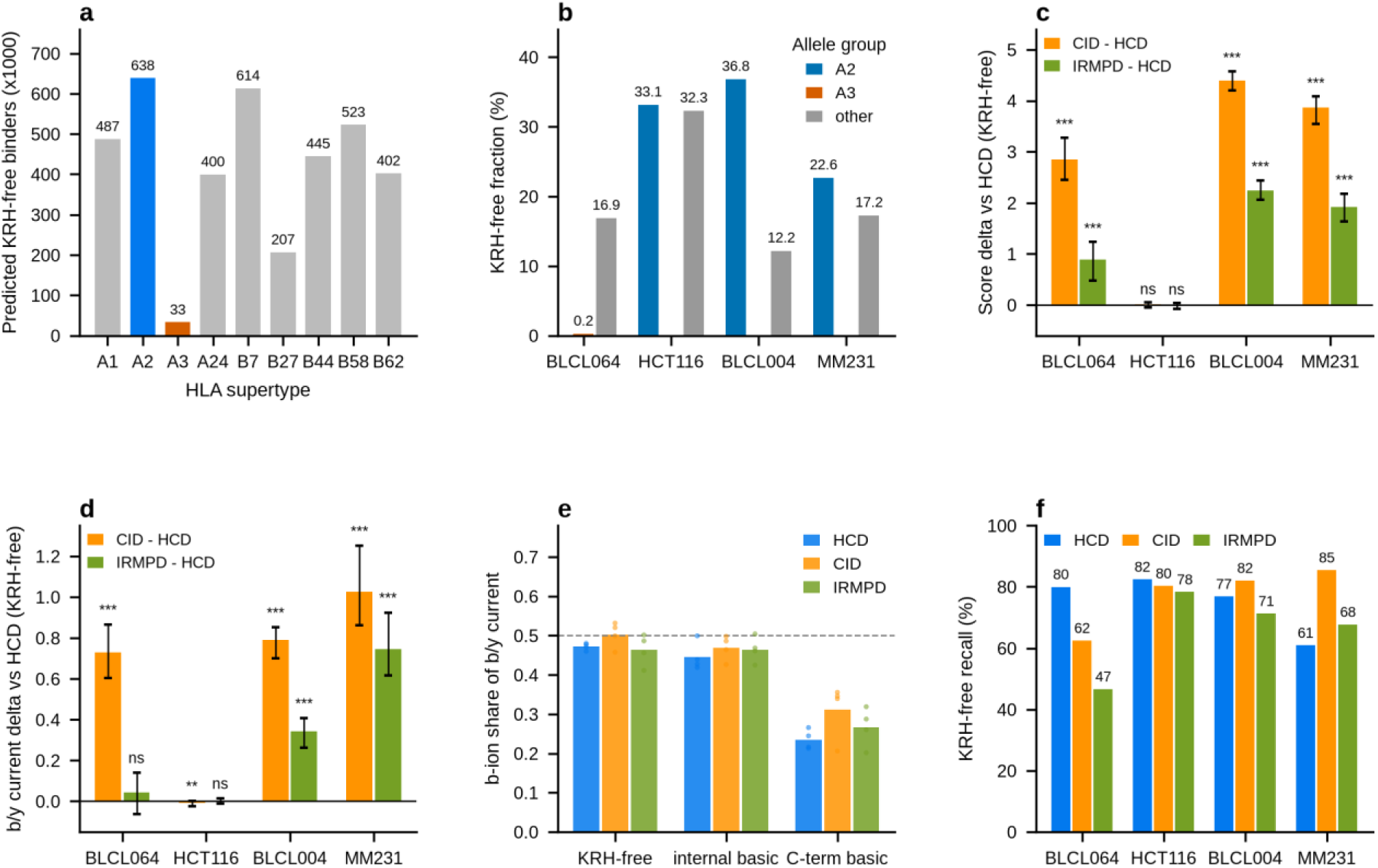
KRH-free peptides link HLA supertype chemistry to fragmentation-method performance. (a) Predicted KRH-free binders (MHCflurry rank < 2) per HLA supertype among 13,629,612 unique KRH-free 8-12-mers enumerated from the reviewed human proteome; A2 (blue) and A3 (orange) highlighted. (b) KRH-free fraction of eluted ligands by assigned allele group per cell line (BLCL064 carries no A2-supertype allele). (c,d) Paired per-peptide spectral-quality deltas versus HCD for KRH-free peptides: MSFragger hyperscore (c) and summed b/y ion current (d); bars, median delta; whiskers, 95% bootstrap CI; stars, Benjamini-Hochberg adjusted Wilcoxon signed-rank p (*** < 0.001, ** < 0.01, ns >= 0.05). (e) b-ion share of total b/y current by residue class and method (bars, mean across cell lines; points, individual cell lines; dashed line, equal b/y split). (f) Per-method recovery of the combined KRH-free peptide repertoire per cell line. The combined repertoire is all KRH-free peptides identified by at least one of the three methods in that cell line; each bar shows the percentage of that repertoire recovered by the individual method. Because methods share peptides, the three bars sum to more than 100%.

Because they lack a sequestering basic residue, KRH-free peptides fragment under the mobile-proton regime. Consistent with this, their b- and y-ion series carried nearly equal current in all three methods (b-share 0.46–0.53). In contrast, C-terminal-basic peptides were strongly y-dominant (b-share 0.20–0.36) regardless of method (**Figure 12e**). The y-ion bias of CID spectra is therefore a property of C-terminal-basic peptides rather than of the fragmentation method itself.

For KRH-free peptides, CID and IRMPD produced better spectra than HCD of the same peptide. Spectral quality here is defined by two quantities measured on the same peptide acquired by two methods: the MSFragger hyperscore, which reflects how many b and y ions match and how intense they are, and the summed b/y ion current, that is the sum of the base-peak-normalised intensities of all matched b and y ions, which reflects how much of the total signal ends up in assignable fragments. Peptides were paired by sequence, one best charge 2+ PSM per method, so that precursor charge and sequence are held constant and only the activation method differs. Paired per-peptide comparisons showed higher hyperscores (median CID–HCD delta +2.8 to +4.4 across BLCL064, BLCL004 and MM231; rank-biserial r 0.79–0.97; Benjamini–Hochberg FDR < 0.001; sign-consistent in all three replicates) and higher summed b/y ion current (CID–HCD +0.73 to +1.03; IRMPD–HCD +0.34 to +0.75 where significant) (**Figure 12c,d; Supplementary Table S11; Supplementary Figure S19**). Consistent with the intensity-concentration mechanism, the background fraction also decreased for CID and IRMPD, whereas matched ions per residue decreased for IRMPD and cosine similarity to the predicted spectrum decreased marginally - both expected when signal is concentrated into fewer fragments and when the reference model was trained on HCD spectra, respectively (**Supplementary Figure S19**). Representative mirror spectra of the two KRH-free 9mers with the largest paired advantage in each cell line illustrate this directly, for instance SVDVTNTTF in BLCL004 (CID score 14.1 versus HCD 2.1; summed b/y current 2.30 versus 0.71) (**Figure 13**). Notably, the advantage scaled with basic-residue content: it was largest for KRH-free peptides, intermediate for internal-basic peptides (+1.7 to +2.4), and smallest for C-terminal-basic peptides. Interestingly, the advantage was absent in HCT116 for every class (all median deltas within ±0.09), indicating an acquisition-level rather than peptide-level origin in that line. A parallel look at the three cryptic KRH-free peptides identified by HCD and at least one slow-heating method showed the same direction in two of three cases; the exception, VTLPVQQAW, favoured HCD on search score but still showed greater b/y-current concentration under IRMPD (**Figure 14**). Consistent with the repertoire-level bias, A2-assigned ligands, which are enriched for KRH-free sequences, recapitulated the CID advantage (+2.69 in BLCL004, +2.42 in MM231), whereas A3-assigned ligands showed only the smaller basic-class gain (+1.65 in BLCL064).

**Figure 13.**
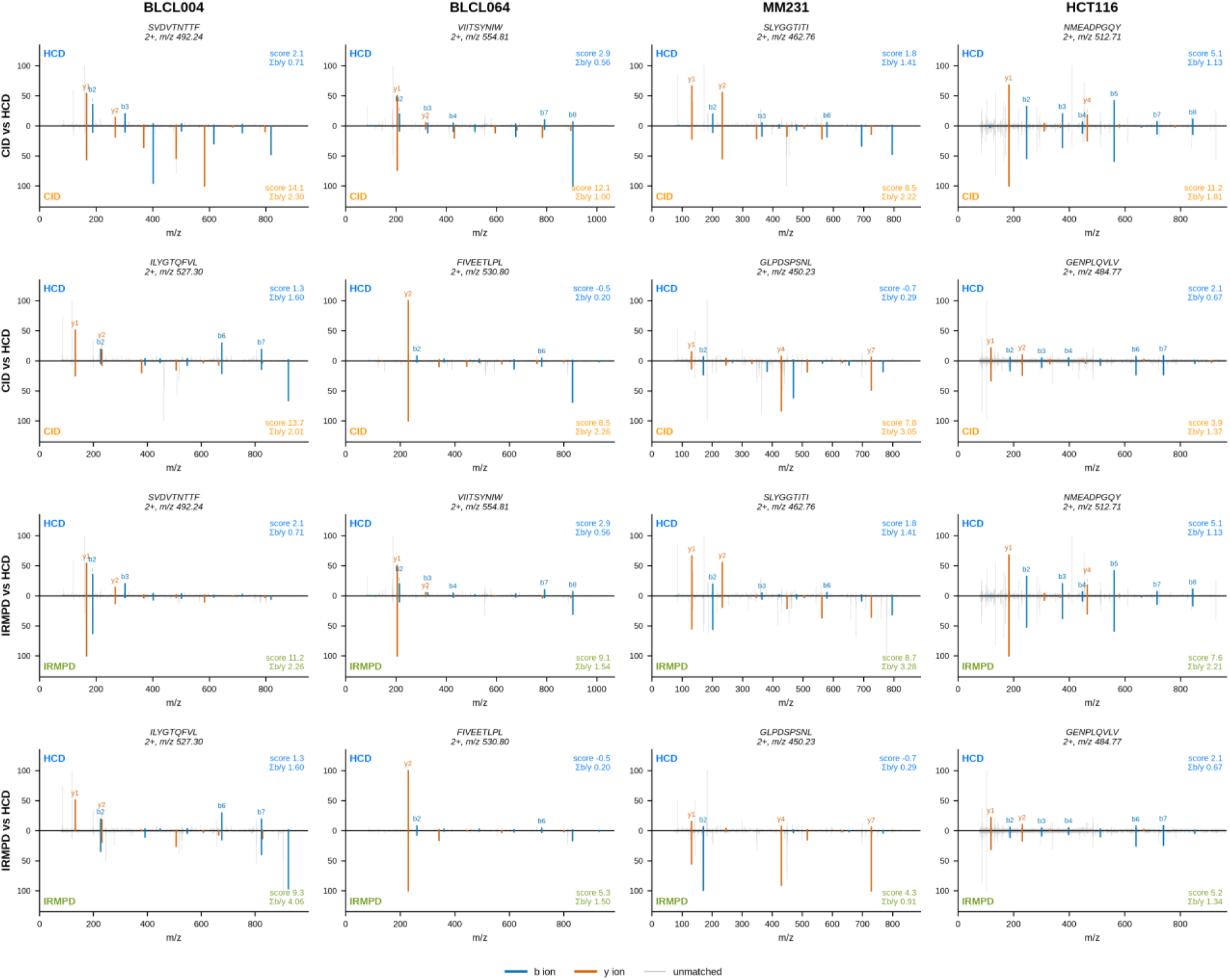
Example KRH-free canonical peptide spectra show the CID/IRMPD spectral-quality advantage. Mirror plots of the best charge-2+ PSM per method for the two KRH-free 9-mers with the largest paired spectral-quality advantage in each cell line (columns); peptide sequence, charge state and precursor m/z are annotated above each panel (all PSMs are cha rge 2 +, so every mirror pair is charge-matched). Rows 1-2, CID (downward) versus HCD (upward); rows 3-4, IRMPD (downward) versus HCD for the same two peptides. Matched b ions (blue) and y ions ( orange) are annotated at theoretical m/z; unmatched peaks in grey. The search score shown is the MSFragger hyperscore; summed b/y ion current ( sum of base-peak-relative intensities of matched ions) is also shown for ea ch spectrum. HCT116 shows no average method difference (acquisition-level flatness); its panels represent the extreme tail of its per-peptide distribution.

**Figure 14.**
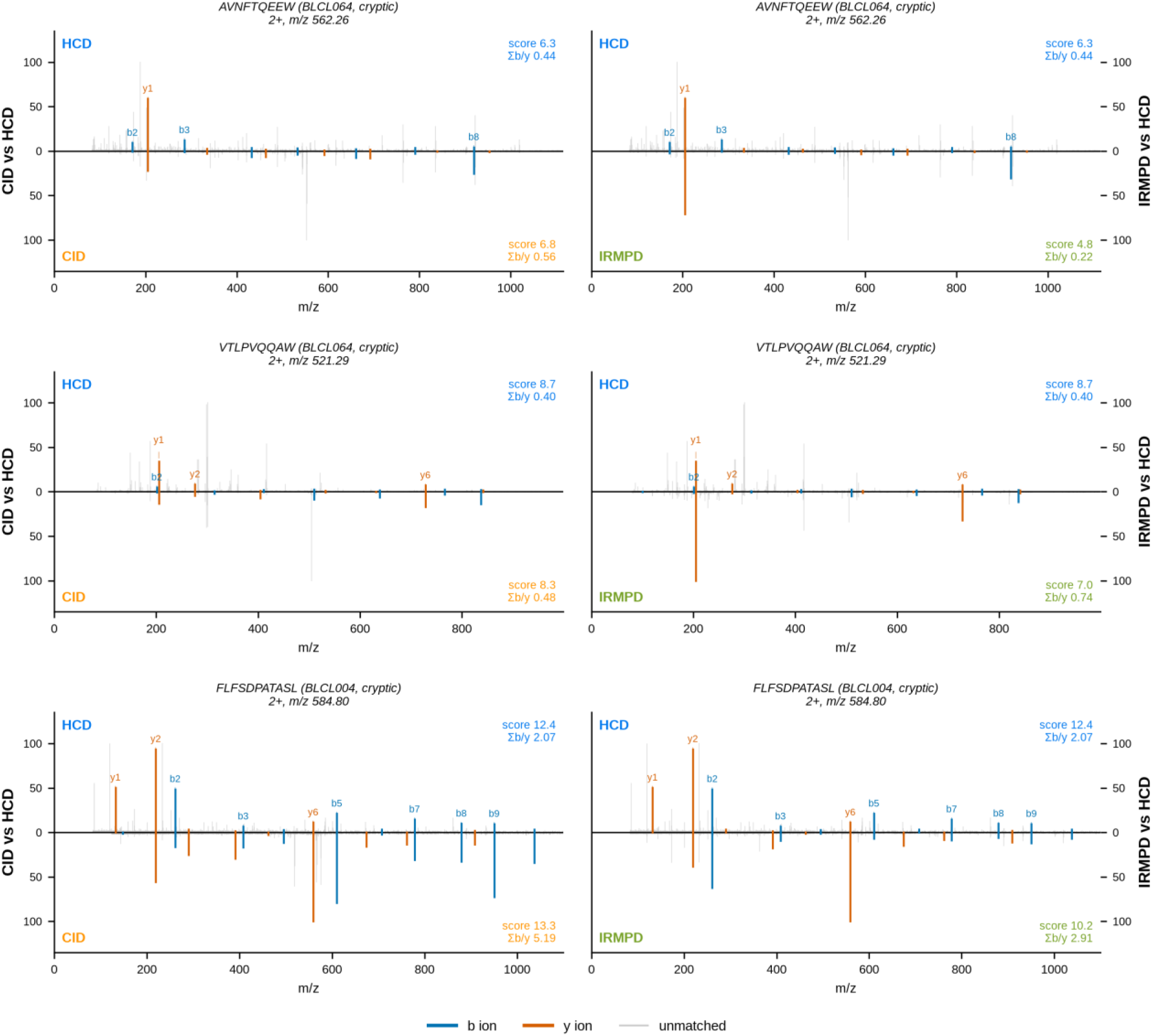
Example KRH- free cryptic peptide spectra. Mirror plots as in Figure 13 for the three cryptic (Ribo-seq ORF-derived) KRH-free peptides identified by HCD and at least one slow-heating method: AVNFTQEEW and VTLPVQQAW (BLCL064) and FLFSDPATASL (BLCL004); peptide sequence, charge state (all 2+) and precursor m/z are annotated above each panel, and the score shown is the MSFragger hyperscore. Left, CID versus HCD; right, IRMPD versus HCD. AVNFTQEEW and FLFSDPATASL show the CID advantage in both score and summed b/y current; VTLPVQQAW modestly favors HCD on score and is shown for completeness, although IRMPD still concentrates more b/y current (0.74 versus 0.40).

This spectral-quality advantage translated into identification-level gains where A2-supertype alleles dominate the repertoire. For example, CID recovered 85.4% of the union KRH-free peptide set in MM231 versus 60.9% for HCD, and 82.1% versus 76.8% in BLCL004. In contrast, HCD recovered more in BLCL064 (80.0% versus 62.5%), the only A2-negative, A3-only line (**Figure 12f**; **Supplementary Figure S20**). This pattern mirrors the allele-level complementarity of **Figure 7c**, where all six MM231 alleles favoured CID and IRMPD while the two A3-supertype alleles A*11:01 and A*33:03 favoured HCD. However, HCD retained the most complete fragment-ion ladders even for KRH-free peptides (charge-matched 2+ completeness 11.7% versus 3.4% CID and 3.0% IRMPD; **Supplementary Figure S21**). The CID and IRMPD advantage is therefore one of fragment-ion intensity concentration rather than bond coverage, a coverage against intensity trade-off that follows from the mobile proton model [52]. Together with the allelic analysis, these results indicate that fragmentation-method choice can be matched to sample HLA composition: CID or IRMPD for A2-supertype-dominated samples, and HCD where basic-anchor alleles dominate.

## Discussion

This study set out to test whether activation chemistry other than HCD has anything to offer HLA class I immunopeptidomics. It does, and the benefit takes two forms: peptides that HCD does not return at all, and better spectra of peptides that both methods return. Both follow from the same underlying trade-off between two desirable properties of a fragmentation method. HCD distributes detectable fragment-ion signal across a more complete backbone-ion ladder, whereas CID and IRMPD can concentrate a larger fraction of the observed signal into fewer matched fragments, yielding stronger individual ions and higher search scores for the same peptides (**Figure 6**, **Figure 12c,d**). Which side of this trade-off benefits a peptide depends on its basic-residue content. Peptides without lysine, arginine or histidine, the three basic amino acids, fragment under the mobile proton model [52], splitting ion current nearly evenly between b- and y-ions in all three methods (b-share 0.46–0.53). Peptides with a basic residue at the C terminus, by contrast, trap the proton there and produce mostly y-ions regardless of method (b-share 0.20–0.36) (**Figure 12e**). The y-ion bias often attributed to CID is therefore a property of the peptide, not the method. Accordingly, the CID/IRMPD spectral-quality advantage is largest for peptides lacking basic residues (median score improvement +2.8 to +4.4 over HCD) and shrinks as basic-residue content increases.

The complementarity observed here operates at three levels. First, at the spectral level, the fragmentation chemistries differ in how they distribute sequence information, with score consequences that track peptide basicity **(Figure 6**, **Figure 12c–e**). Second, these spectral differences translate into partially distinct peptide and allele repertoires, such that each method recovers ligands the others miss. Third, this non-redundancy has a practical consequence: at a fixed acquisition budget, distributing runs across fragmentation methods increases the number of unique HLA ligands recovered and expands the set of novel antigen candidates. The value of CID and IRMPD thus lies not in replacing HCD, but in extending the immunopeptidome accessible to an HCD-only workflow.

This same basicity gradient explains the allele-level differences we observed. HLA alleles whose binding preference includes a basic residue at the C-terminal anchor position were detected more efficiently by HCD, namely A*11:01 (88% lysine at the anchor, −11.2 and −10.4 percentage points for CID and IRMPD) and A*33:03 (94% arginine), both A3 supertype members. Conversely, all six alleles in MM231, none of which requires a basic anchor, favored CID/IRMPD (+3.2 to +4.4 pp), and across all 21 alleles tested the detection-share difference tracked C-terminal anchor basicity (**Figure 7c,g**). The KRH-free analysis extends this from individual alleles to the whole repertoire. The A2 supertype, whose preferred C-terminal anchor is hydrophobic rather than basic, tolerates KRH-free peptides: 23–37% of A2-assigned ligands were KRH-free. In A2-positive cell lines, CID recovered up to 24.5 pp more of the KRH-free repertoire than HCD, whereas in BLCL064, the only line carrying A3-supertype alleles, which require a basic C-terminal anchor, HCD recovered more (Figure 12b,f). Because A2 is among the most common HLA supertypes worldwide, this has broad practical relevance. Fragmentation method can be matched to sample HLA composition: CID or IRMPD when A2-supertype alleles dominate, HCD when basic-anchor alleles dominate. When composition is unknown or mixed, using more than one method is complementary, not redundant.

The spectral-quality differences also provide context for the rescoring results. CID and IRMPD spectra of KRH-free peptides reach the search engine with higher intrinsic confidence, yet fine-tuned rescoring models added genuine identifications on top of method effects for all three methods (**Figure 2**). Fragmentation choice and rescoring thus act on different levels: one changes the spectrum itself, the other improves the confidence assigned to it. Whether rescoring models capture method-specific fragmentation patterns differently for KRH-free and for basic peptides, and whether the benefit of rescoring therefore differs by method and peptide class, remains open, and the per-method models generated here are a starting point for addressing it.

The larger rescoring gains observed for the cryptic peptides (median +54.5% versus +17.1% for the standard database; **Figure 2b**) reflect the small cryptic baseline rather than a biological difference in peptide behavior. Although the cryptic database contains more sequences than the reviewed Swiss-Prot proteome (23,230 non-canonical open reading frames versus 20,431 entries), these ORFs are far shorter (median 26 versus 559 residues), so the candidate search space is approximately 20-fold smaller (2,583,860 versus 52,221,465 distinct 8-12mers) and does not inflate the FDR threshold. The cryptic identified repertoires are instead 1–2 orders of magnitude smaller than their standard-database counterparts, so rescuing even a modest number of additional peptides yields a large percentage gain relative to the small baseline. The four cell line × method combinations with the smallest cryptic yields (12 to 29 peptides before rescoring) showed no net change, consistent with a denominator too small to absorb additional identifications.

What this work establishes is potential rather than a finished method. CID and IRMPD return ligands that HCD does not, and better spectra for a chemically defined subset of the repertoire, on an instrument where all three are available in a single run schedule. Turning that into routine gain now requires building the infrastructure these chemistries currently lack. Three things are needed. First, fragment-intensity and retention-time models trained on CID and IRMPD immunopeptidome data rather than fine-tuned from HCD, since prediction quality sets the ceiling on rescoring and, in turn, on how much of the alternative-method repertoire is recoverable. Second, acquisition schemes that allocate activation method by precursor properties, so that a peptide is fragmented by the chemistry most likely to yield an interpretable spectrum instead of by a fixed instrument setting. Third, evaluation in data independent acquisition, where the lower chemical background of the two slow-heating methods may improve deconvolution of the highly multiplexed spectra that limit DIA immunopeptidomics [14,53]. Until those pieces exist, the practical recommendation from this work is narrow and usable: where instrument time allows more than one run per sample, spend it on more than one fragmentation method, and where the HLA type is known, weight the allocation towards CID or IRMPD for A2-supertype-dominated samples and towards HCD where A3-supertype alleles dominate.

## Data availability

The mass spectrometry proteomics data, including raw files, search results and the parameter files used for MSFragger, AlphaPeptDeep fine-tuning and Percolator rescoring, have been deposited to the MassIVE repository and are available at https://doi.org/10.25345/C5N58D150. Username for web access: MSV000103328_reviewer; Password: Bjjc6FYcf7L1pOp7

## Author contributions

T.L.K.S., F.S., N.S., and T.S. conducted the experimental work and acquired the data. T.L.K.S., F.S, P.F., and G.G. analysed the data. T.L.K.S., P.F., R.B.S., F.S., C.M., and J.H. designed the research studies. T.L.K.S., F.S., and P.F. wrote the manuscript. T.L.K.S., R.B.S., F.S., C.M., and P.F. edited the manuscript. P.F. acquired funding and supervised the project. All authors have read and agreed to the published version of the manuscript.

## Acknowledgements

We acknowledge A/Prof Joshua Ooi for the provision of B-lymphoblastoid cell lines (B-LCLs). All proteomic analyses were conducted at the Clinical Proteomics Node (CPN) of the Monash Proteomics and Metabo-lomics Platform (MPMP), where infrastructure enabled by Bioplatforms Australia (BPA) and the National Collaborative Research Infrastructure Strategy (NCRIS) was utilised. This work was further supported by Monash eResearch capabilities, including Research Data Storage and Nectar Research Cloud.

## Supplementary material

**Supplementary Figure S1.**
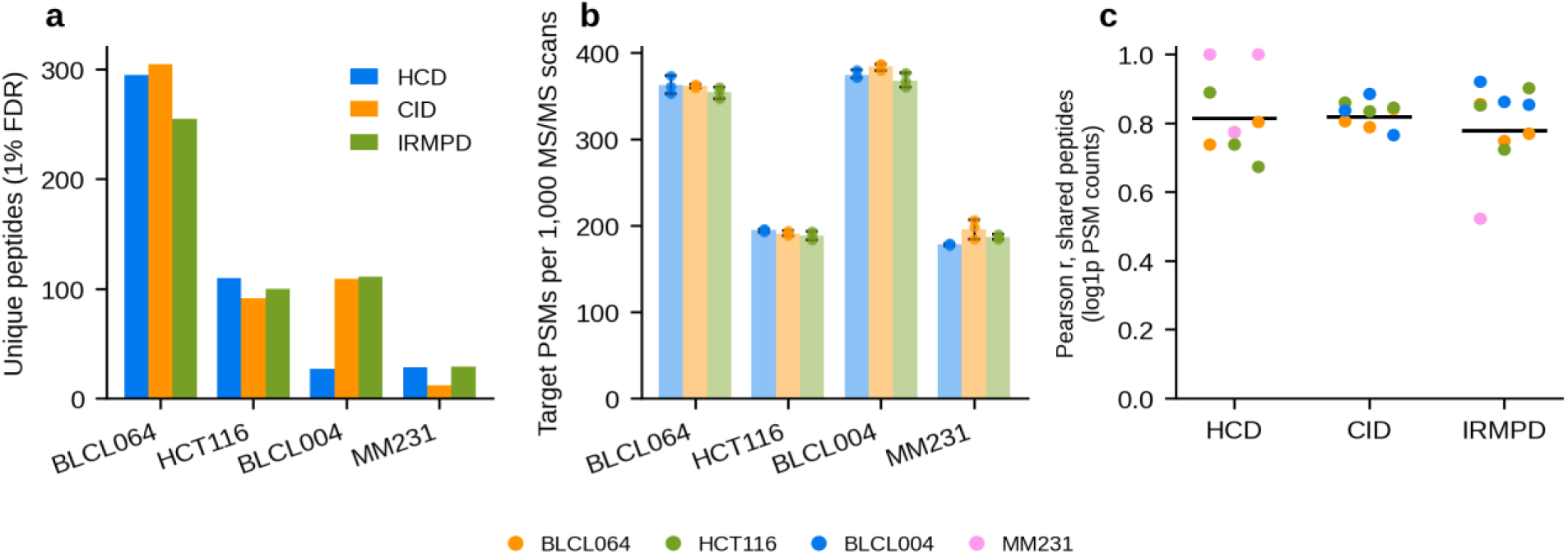
Identification quality control, cryptic database. Panels as in Figure 3, for the four HLA-typed cell lines. Cryptic repertoires are 1-2 orders of magnitude smaller than standard-database repertoires; pairwise Pearson correlations are computed on small shared peptide sets (down to <10 peptides for the lowest-yield combinations) and are correspondingly noisy; combinations with fewer than five shared peptides are omitted.

**Supplementary Figure S2.**
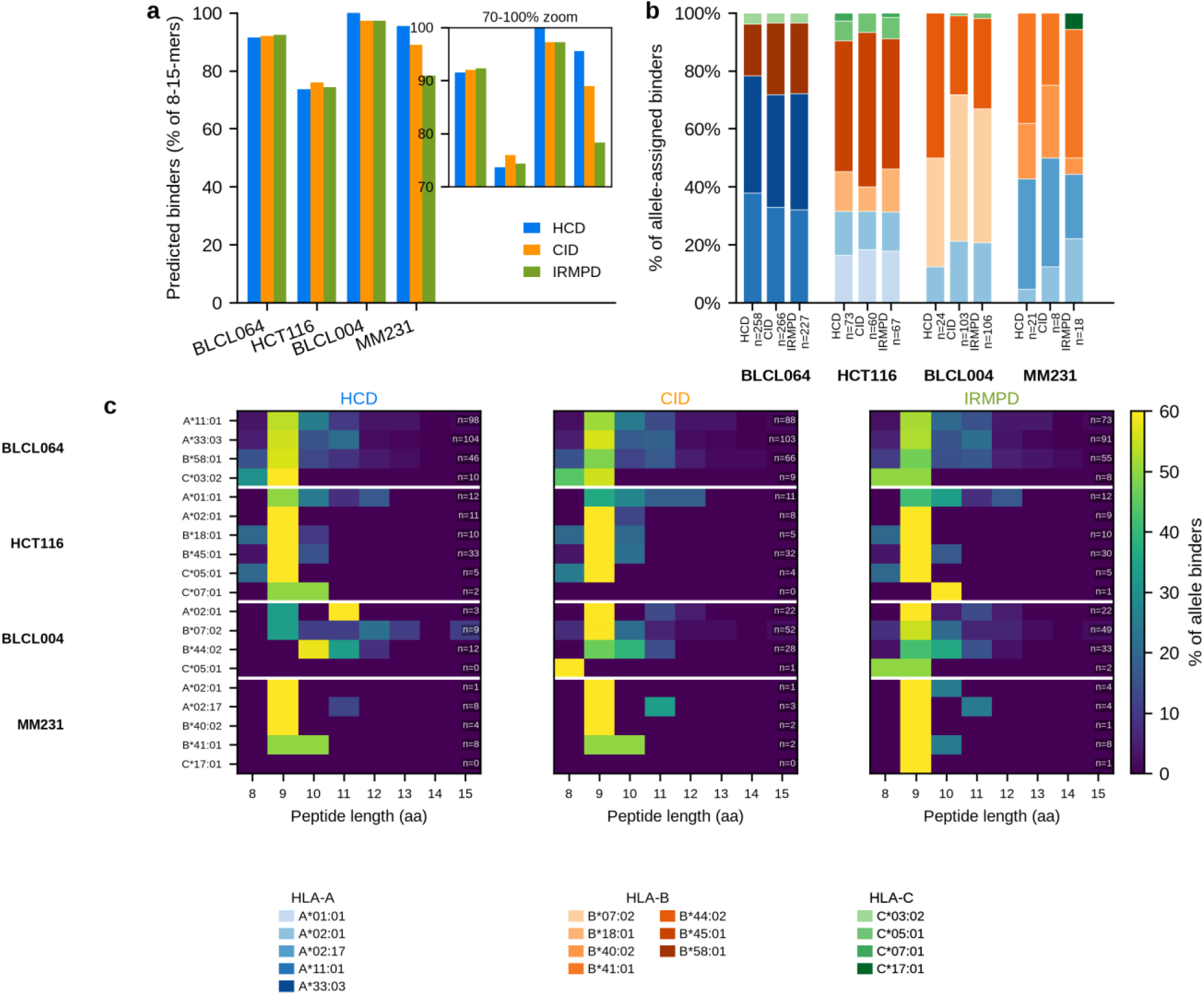
HLA-binding quality control, cryptic database. Panels as in Figure 4a-c; panel c shows per-method length distributions as in Figure 4c, with the number of binders (n) annotated per allele and method because cryptic repertoires are small. Binder fractions of cryptic peptides remain high (73.7-100%), supporting their identity as genuine HLA ligands.

**Supplementary Figure S3.**
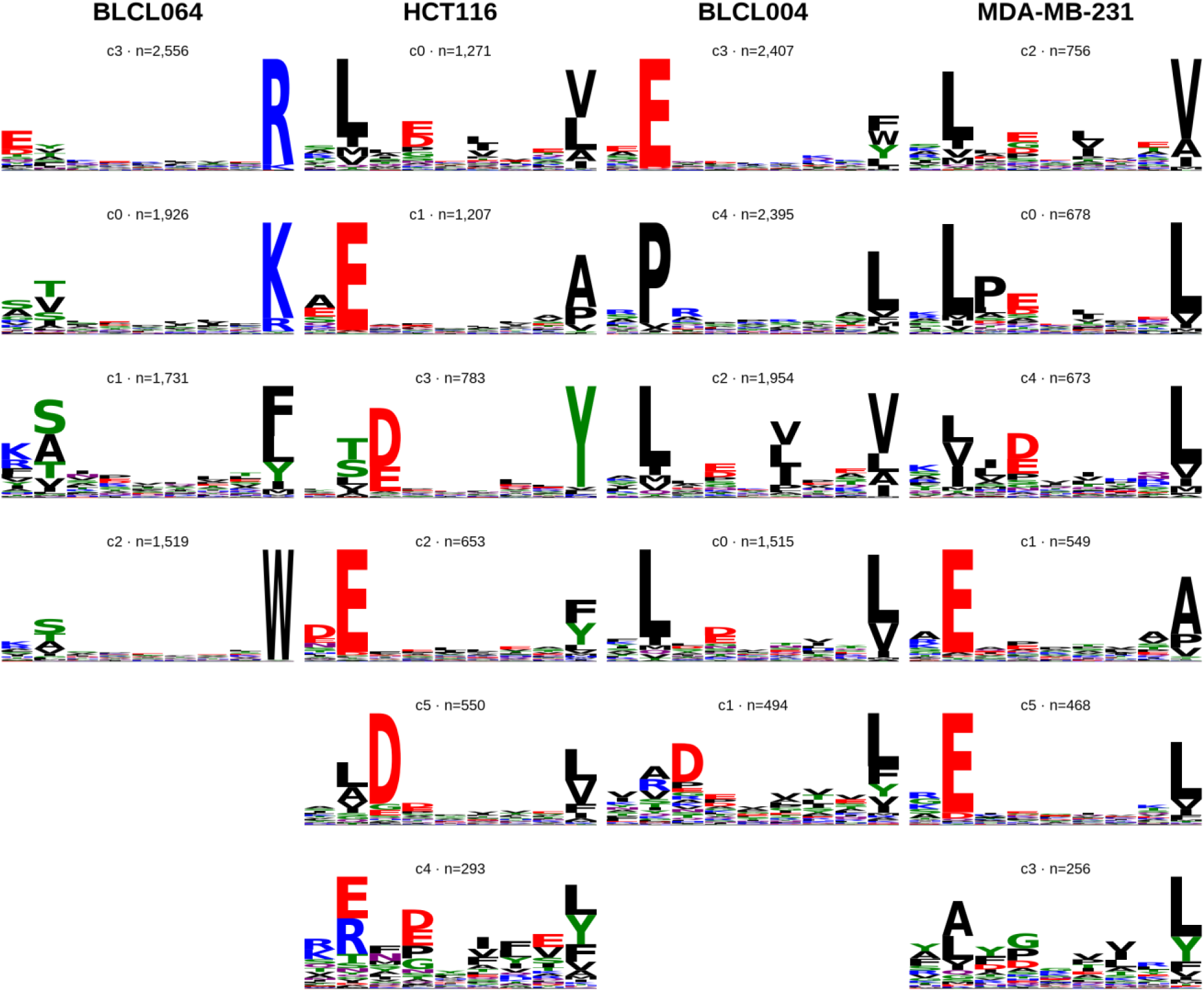
Union GibbsCluster 2.0 deconvolution of the 9mer repertoires. As Figure 5, but clustering the union of unique 9mers across fragmentation methods per cell line ( K = number of typed alleles; best of three seeds by final KLD). Columns = cell lines; rows = clusters ordered by descending size; captions give the native cluster ID and cluster size (n).

**Supplementary Figure S4.**
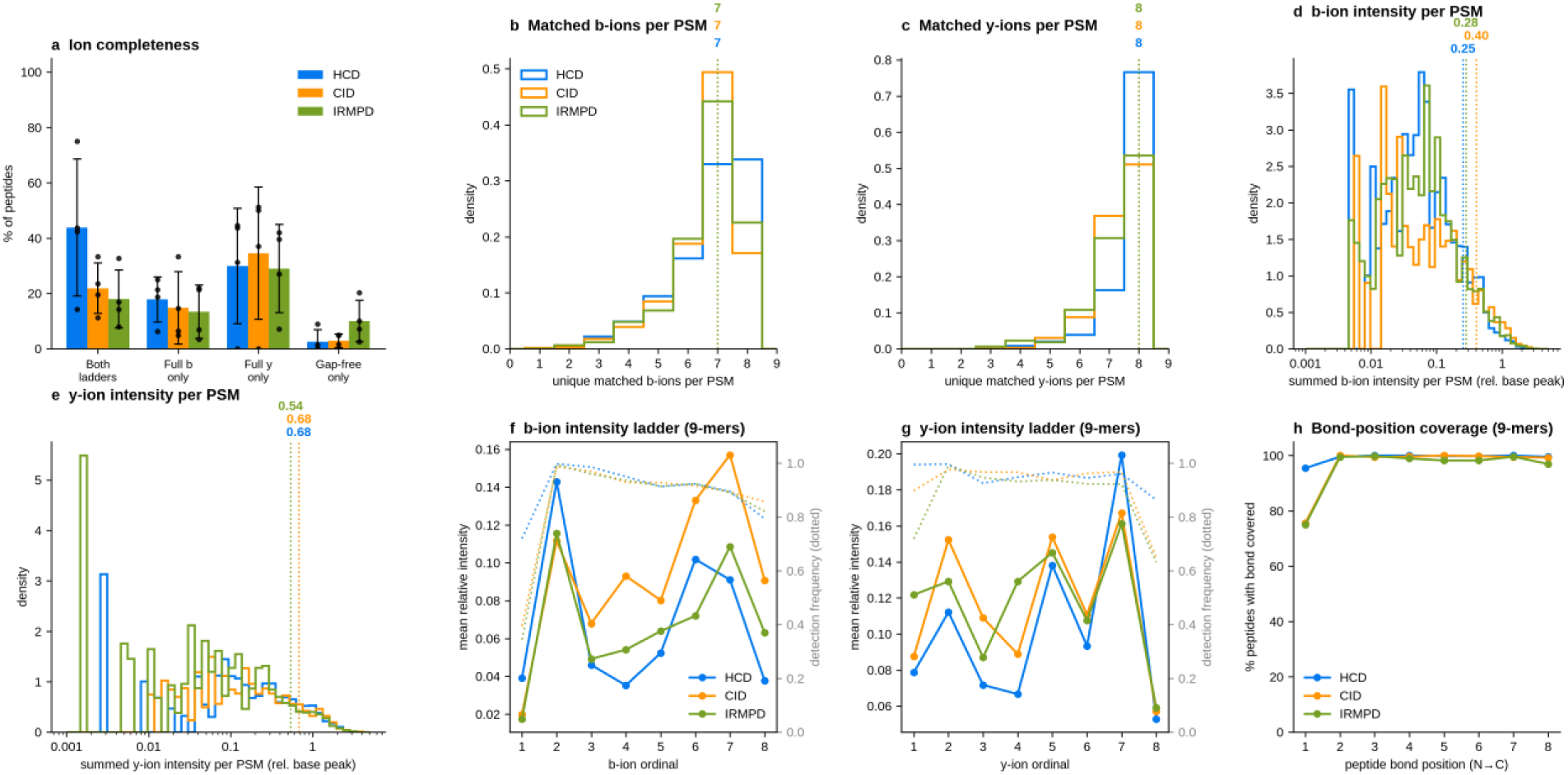
Fragment-ion coverage and intensity balance, cryptic database. Panels as in Figure 6, for the four HLA-typed cell lines, restricted to the 9mer predicted binders (best EL %Rank < 2). Cryptic 9mer-binder repertoires are small (4-144 peptides per cell line x method), so per-cell values are noisier; the same qualitative trends hold (HCD more complete ladders; CID/IRMPD shifted toward b-ion intensity).

**Supplementary Figure S5.**
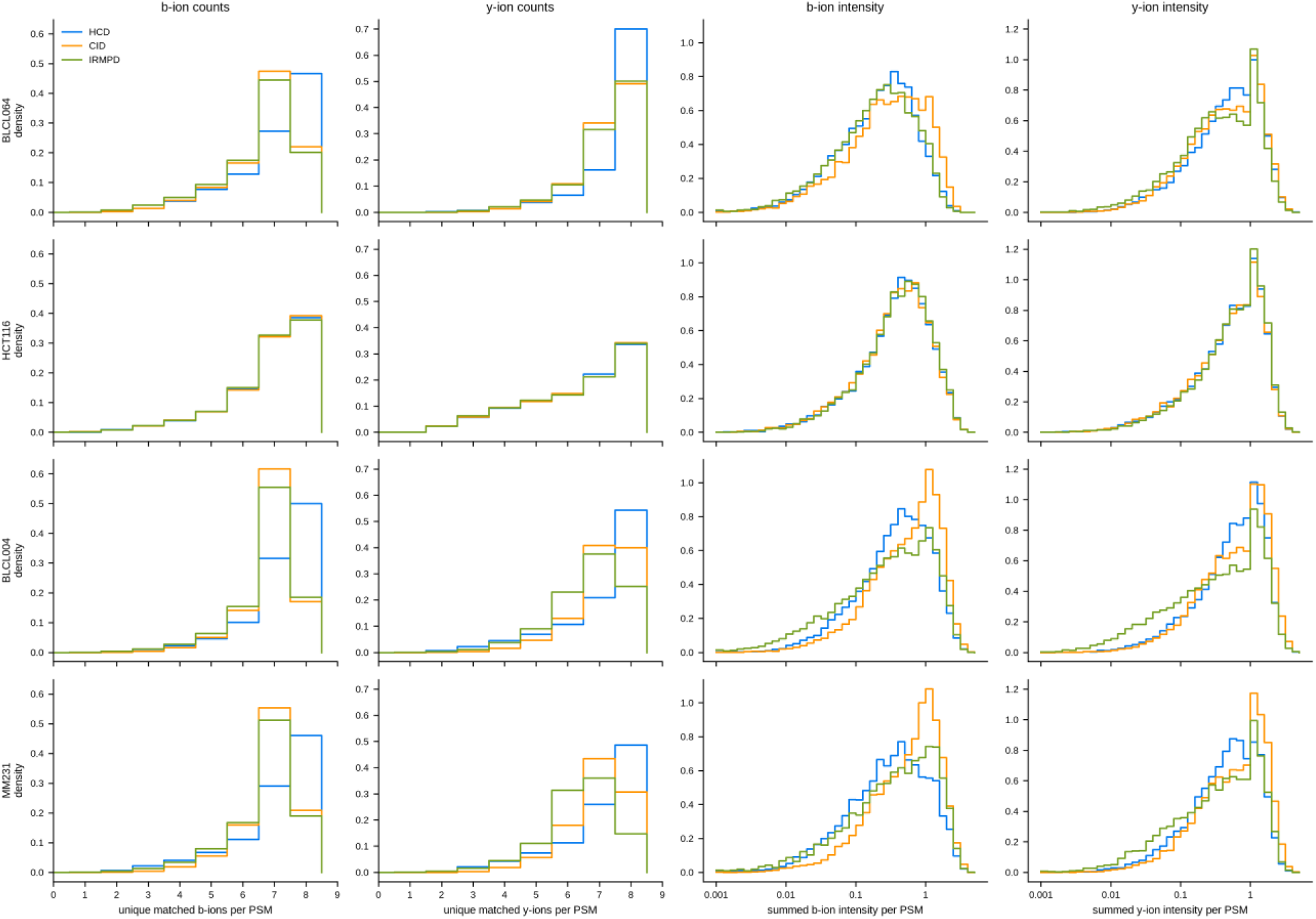
Per-cell-line distributions of matched b- and y-ions per PSM (standard database), restricted to the 9mer predicted binders (best EL %Rank < 2) underlying Figure 6. **Density-normalised step histograms of unique matched ions per PSM (distinct fragment ordinals per series; b- and y-series, left two columns)** and summed matched-ion intensity per PSM (right two columns, log10 scale, relative to the base peak), one row per cell line; color = fragmentation method. These per-cell-line distributions underlie the pooled distributions of Figure 6b-e and show whether cross-method balance shifts arise from b-ion gains, y-ion losses, or both. Cryptic per-cell-line distributions are too noisy (down to tens of PSMs per cell line x method); pooled cryptic distributions are in Supplementary Figure S4.

**Supplementary Figure S6.**
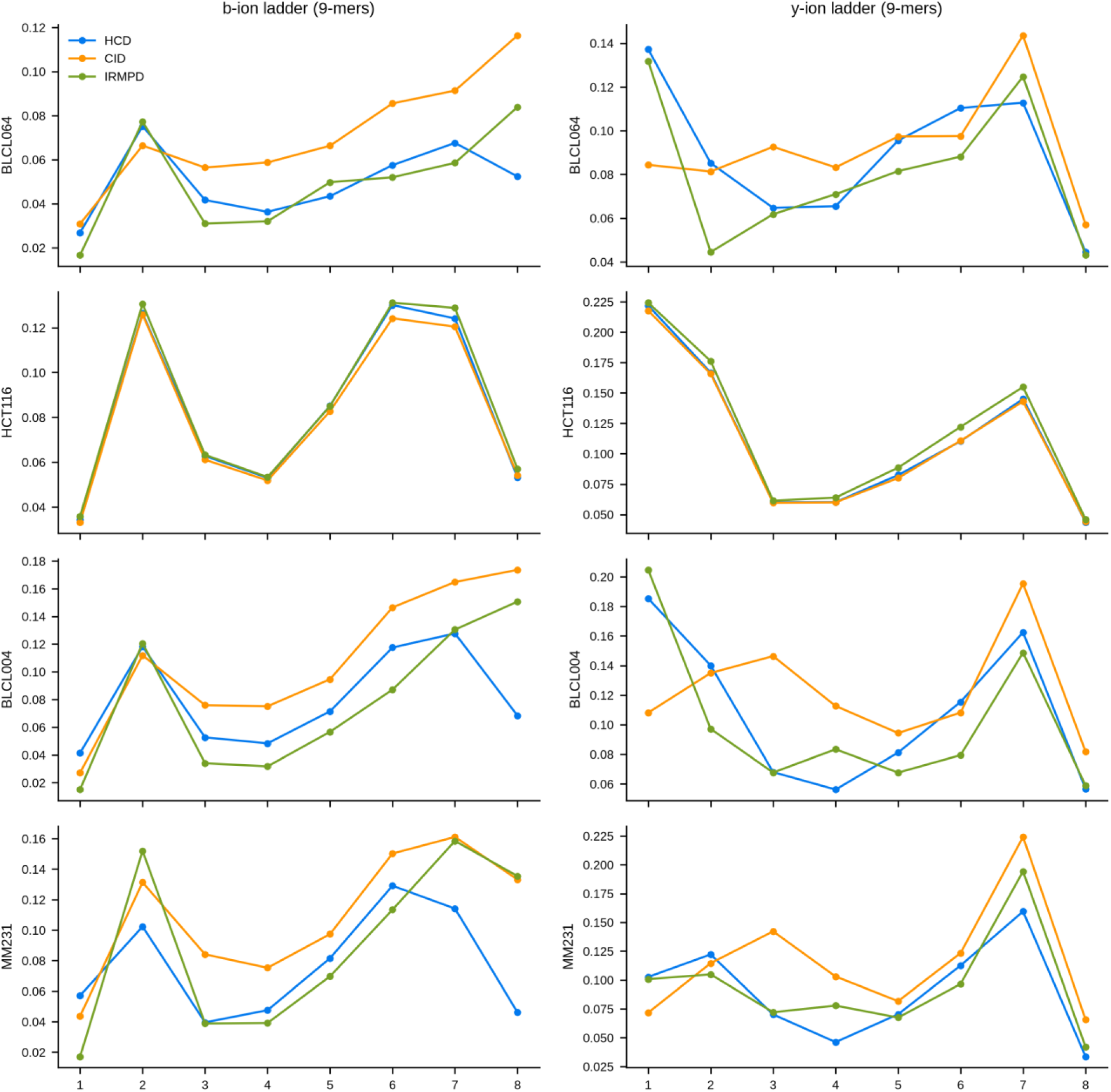
Per-cell-line b- and y-ion intensity ladders of 9mer peptides (standard database). Mean relative intensity per fragment ordinal for each of the four cell lines (rows) and ion series (columns); colors as in Figure 6. The cross-method patterns of Figure 6f,g are reproducible in every cell line.

**Supplementary Figure S7.**
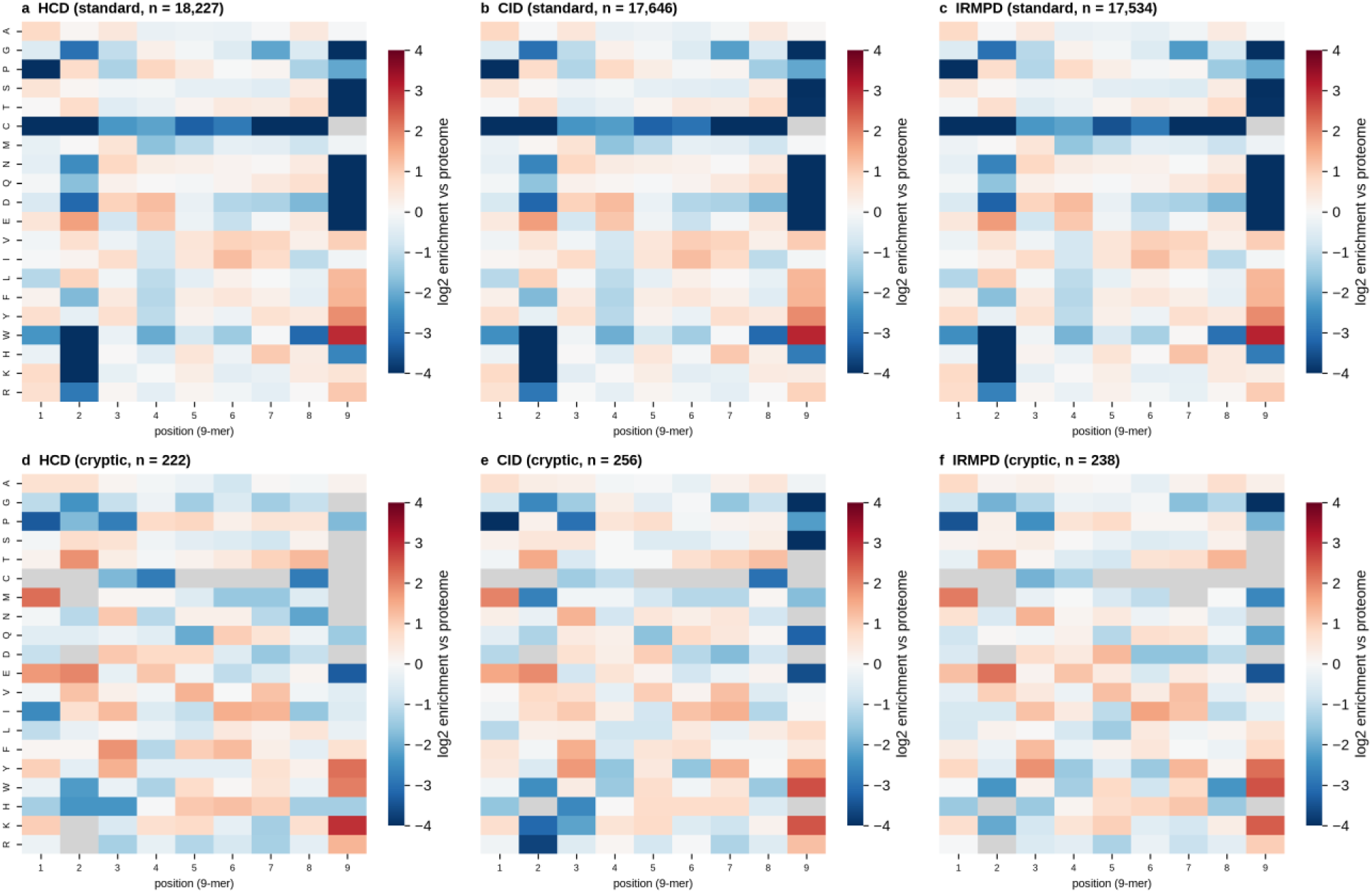
Position-specific amino-acid enrichment of identified 9mers versus the proteome background. log2 ratio of the position-specific amino-acid frequency of pooled identified 9mers (union across the four cell lines) to the corresponding proteome-background frequency, per fragmentation method (columns), for the standard (a-c) and cryptic (d-f) databases. The HLA anchor structure (enriched P2 and C-terminal residues, depleted central positions) is visible in every method; grey cells are amino acids never observed at that position. n = pooled 9mer count.

**Supplementary Figure S8.**
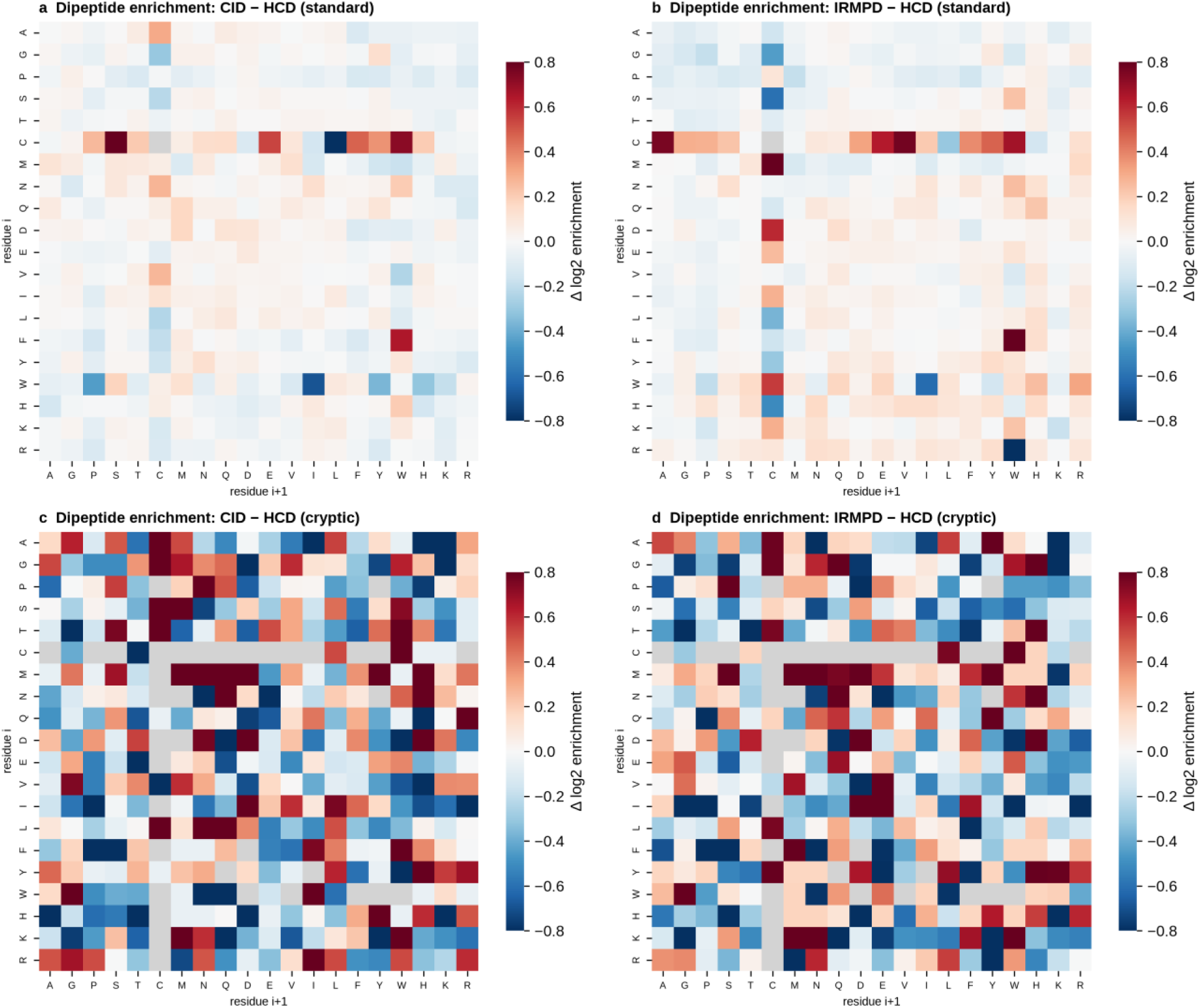
Dipeptide enrichment differences between fragmentation methods. Dipeptide enrichment (log2 observed / expected versus the searched proteome background) difference between CID and HCD (a,c) and between IRMPD and HCD (b,d), mean across the four HLA-typed cell lines, for the standard (a,b) and cryptic (c,d) databases; red = enriched relative to HCD. Grey cells are dipeptides absent from a method’s repertoire (cryptic repertoires are small, so most cryptic cells are unpopulated).

**Supplementary Figure S9.**
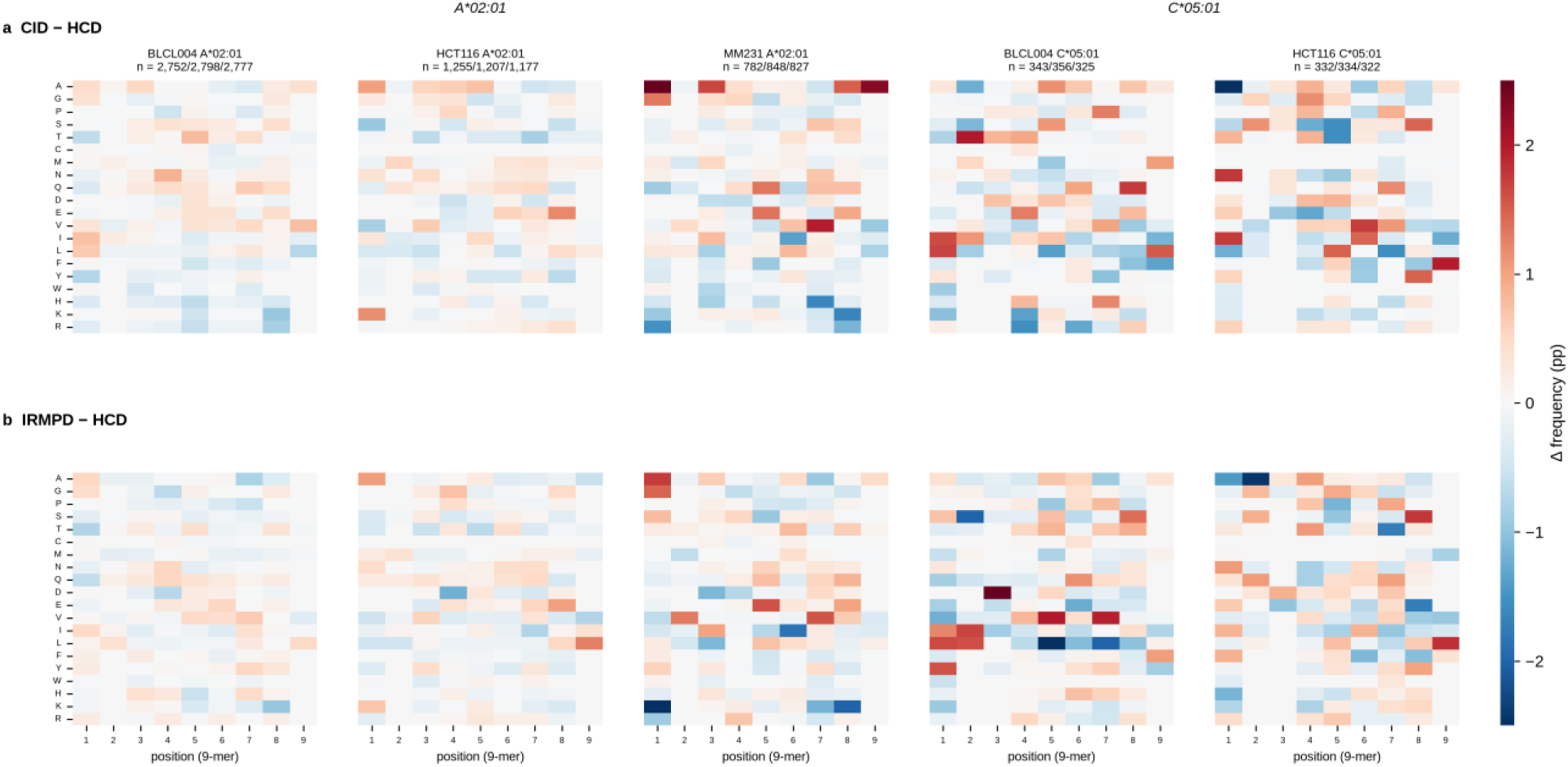
Per-allele position-specific amino-acid differences between fragmentation methods (standard database). Position x amino-acid frequency difference (percentage points), CID minus HCD (a) and IRMPD minus HCD (b), for 9mers assigned to A*02:01 (three cell lines) and C*05:01 (two cell lines) — the alleles with sufficient counts in multiple lines. Within an allele, position-specific frequencies are essentially flat across methods, indicating that fragmentation method shifts allele representation rather than allele-specific motifs. C*05:01 panels are noisier (smaller n). n = per-method 9mer count (HCD/CID/IRMPD).

**Supplementary Figure S10.**
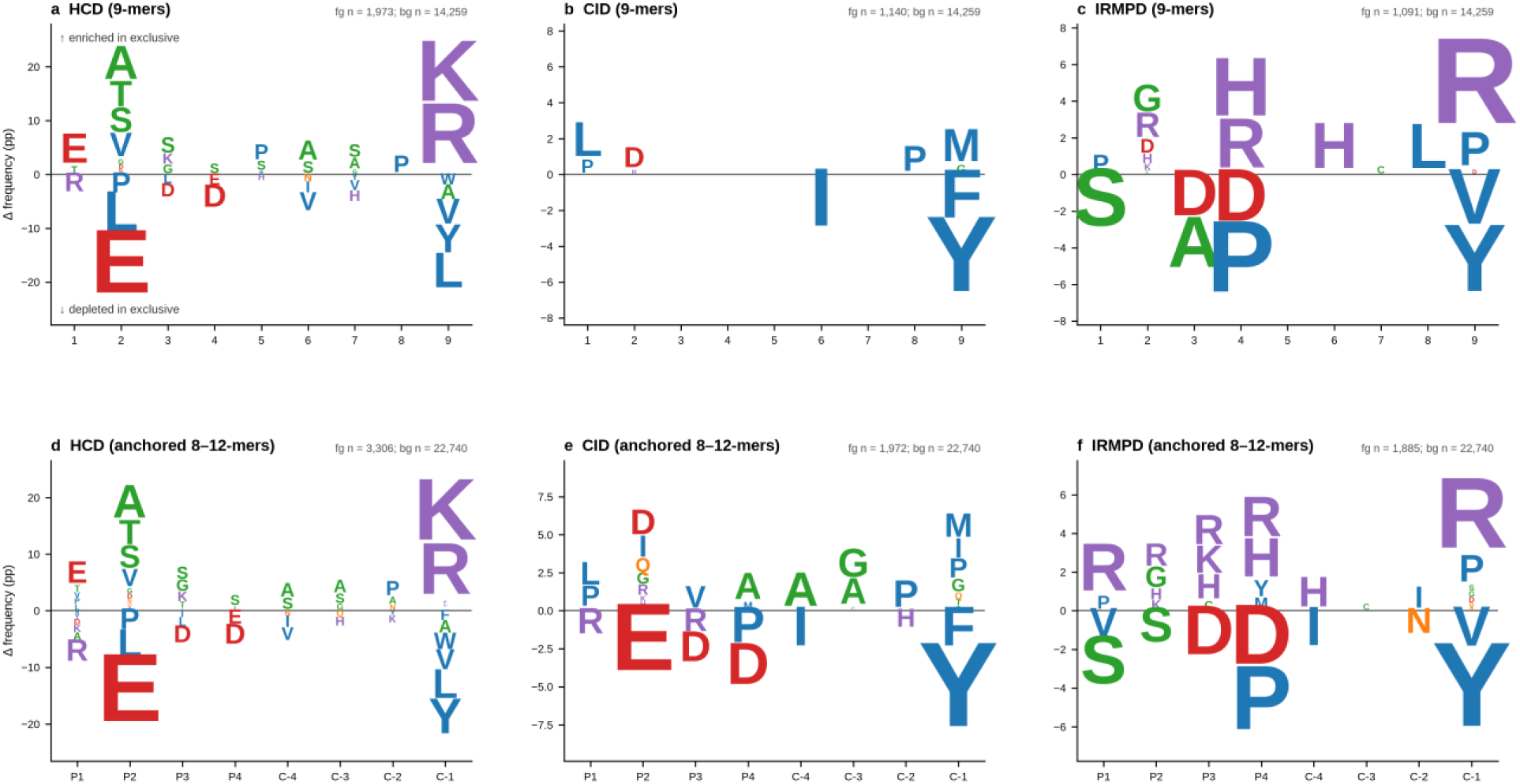
Sequence motifs of method-exclusive peptides (standard database). Two-sample motif comparison of method-exclusive peptides (foreground) versus peptides shared by all three methods (background), pooled across the four cell lines. Letter height = position-specific frequency difference (percentage points); only amino acids significant at Benjamini-Hochberg FDR < 0.05 (two-proportion test) are drawn, stacked upward (enriched in exclusive) or downward (depleted). (a-c) 9mers by position P1-P9; (d-f) 8-12mers aligned by the four N-terminal and four C-terminal residues (P1-P4, C-4 to C-1). fg/bg n = foreground/background peptide counts.

**Supplementary Figure S11.**
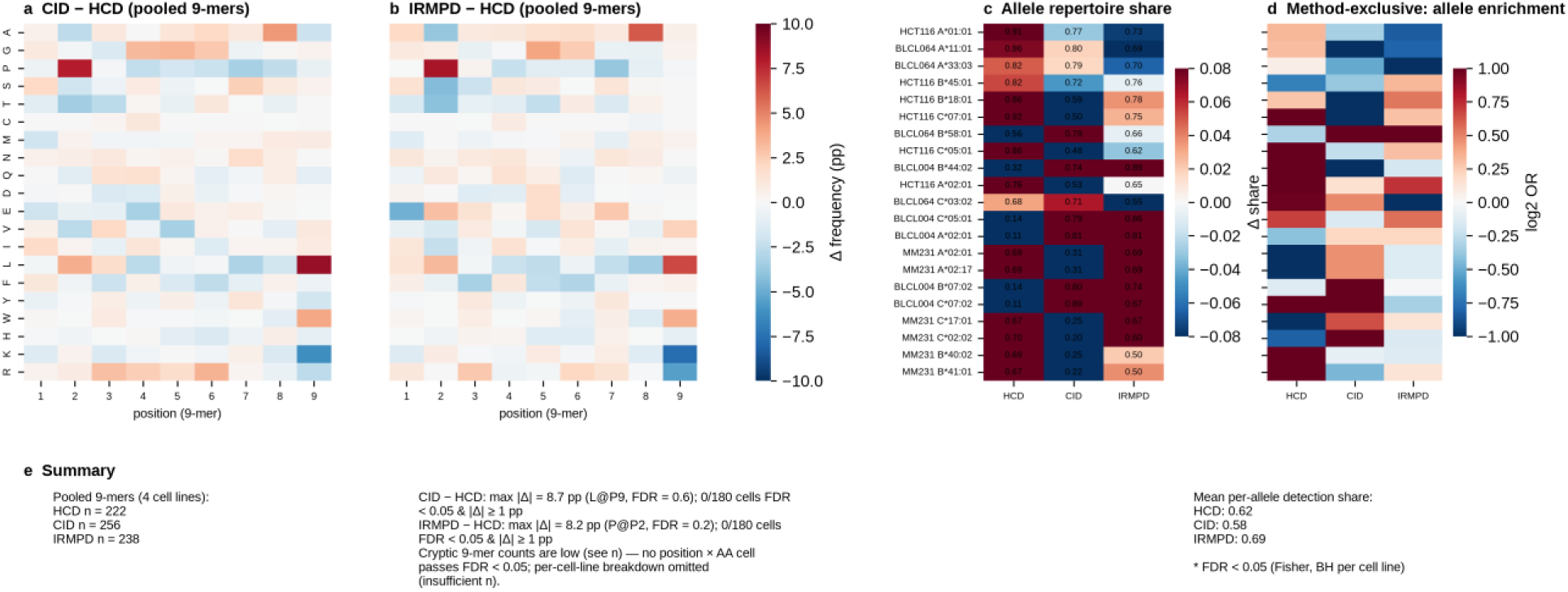
Position-specific amino-acid and allelic preferences, cryptic database. Panels a-d and the summary as in Figure 7 (pooled 9mers only; the per-cell-line breakdown of Figure 7e is omitted because cryptic 9mer counts are too low for stable per-cell position-specific frequencies). No position x amino-acid cell passes FDR < 0.05, and allele-share and enrichment estimates are based on very small cryptic repertoires (12-29 peptides for MM231) and should be read qualitatively.

**Supplementary Figure S12.**
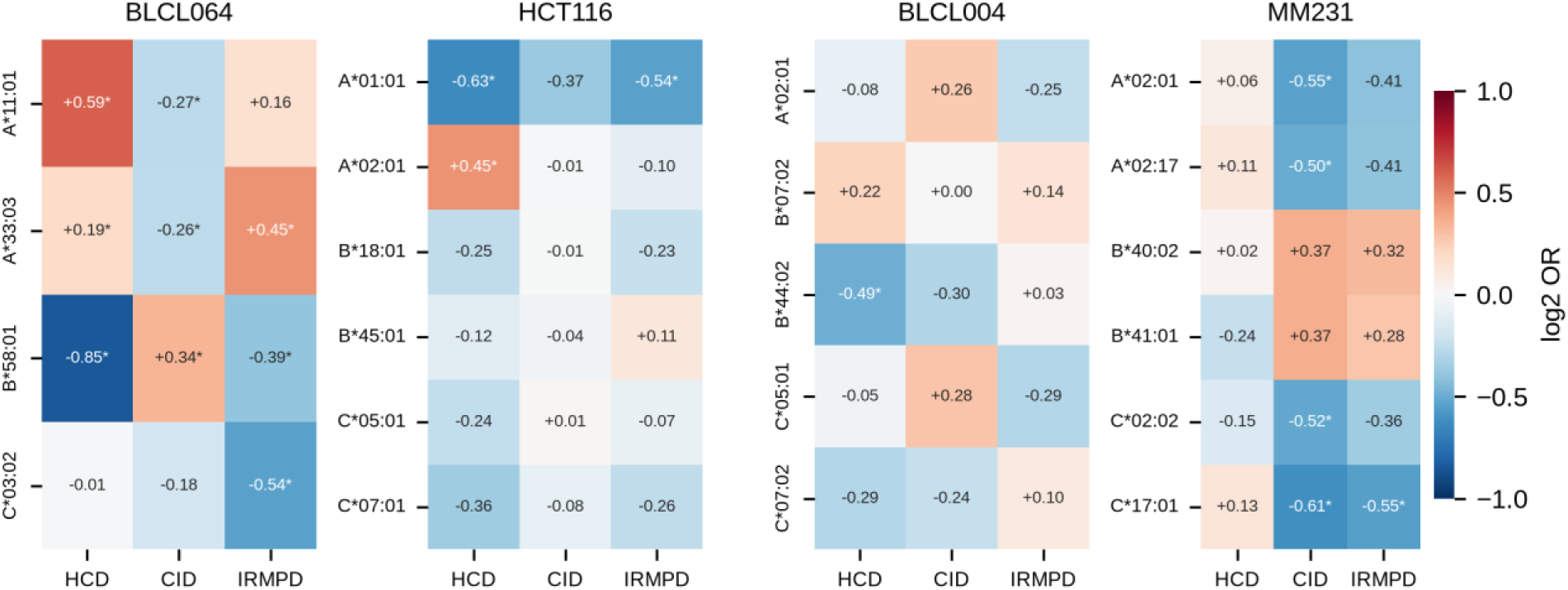
Allelic enrichment of method-exclusive peptides, per cell line (standard database). log2 odds ratio per allele and fragmentation method (Fisher exact test, Benjamini-Hochberg FDR per cell line; * FDR < 0.05) for peptides detected by only one method; values are printed in each cell. Per-cell-line view of the data summarised in Figure 7d.

**Supplementary Figure S13.**
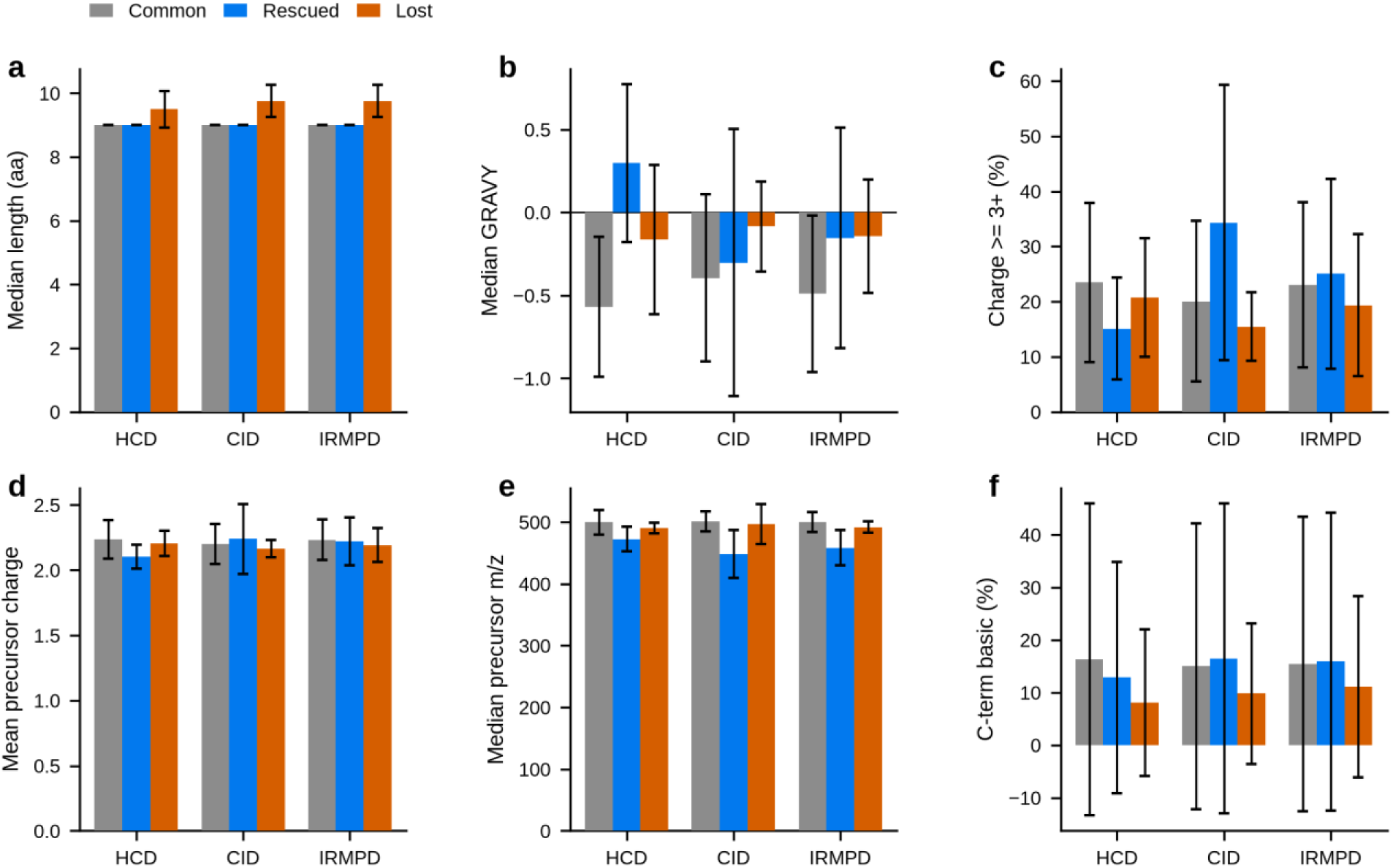
Physicochemical composition of common, rescued and lost peptide sets per method (standard database). Mean +- SD across the four HLA-typed cell lines per set and method: (a) median peptide length, (b) median GRAVY hydrophobicity, (c) fraction of precursors with charge >= 3+, (d) mean precursor charge, (e) median precursor m/z, (f) C-terminal basic-residue fraction. Extends Figure 8d; the underlying per-cell-line values are aggregated in Supplementary Table S5.

**Supplementary Figure S14.**
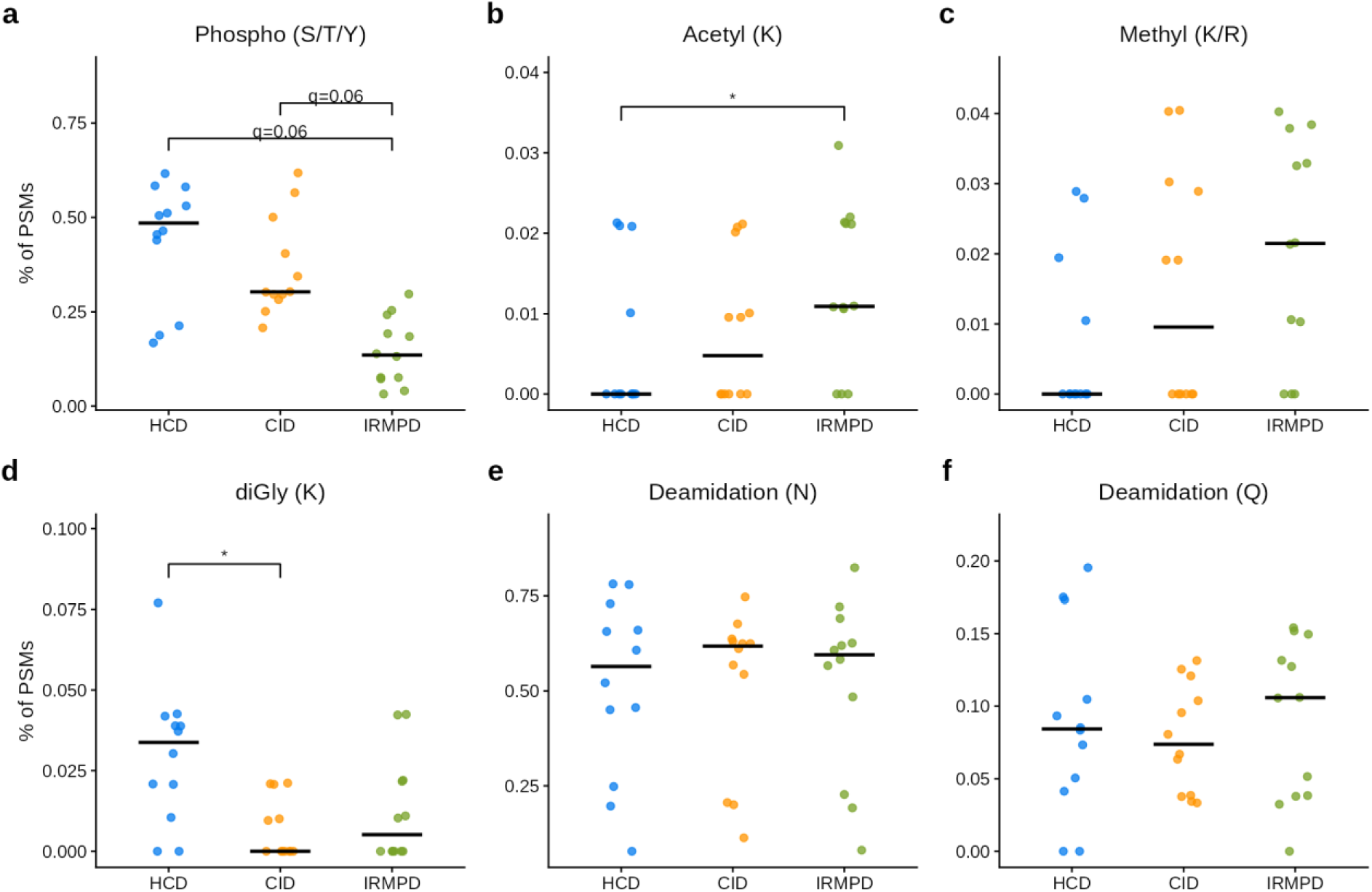
Post-translational-modification rates per fragmentation method (standard database). Open mass-offset search of the pooled 1% FDR PSMs; per-replicate rates (% of PSMs) for the four HLA-typed cell lines (points; black bar = median). Pairwise Wilcoxon tests on per-replicate rates (n = 12 per method) with Benjamini-Hochberg FDR: * q < 0.05; borderline comparisons (q < 0.10) are labeled with the exact q value.

**Supplementary Figure S15.**
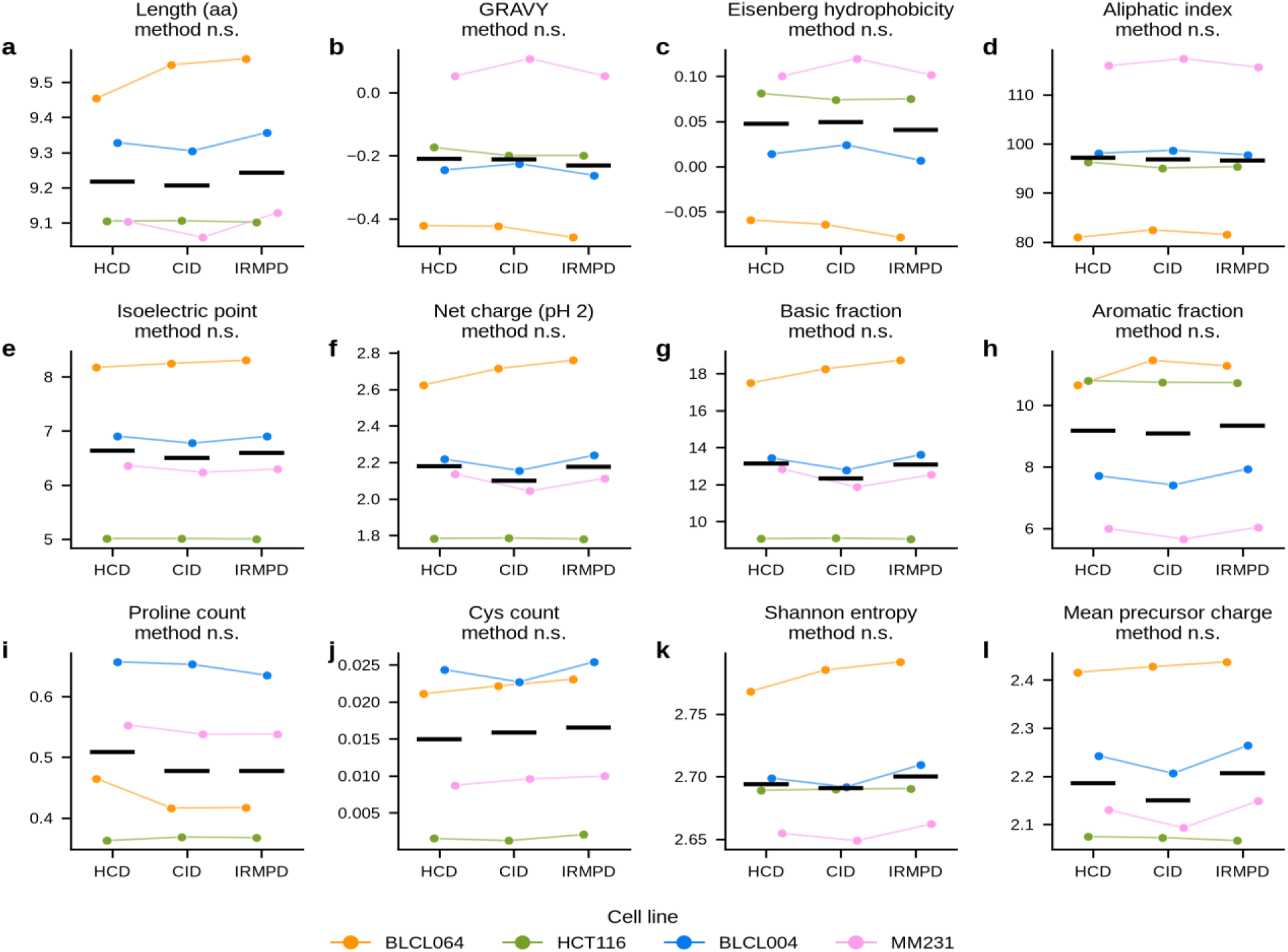
Physicochemical properties of the identified repertoires per fragmentation method, without rescued/lost/common partitioning (standard database). Points = per-cell-line means of unique-peptide sequence properties (a-k) and PSM-level mean precursor charge (l), colored by cell line (connecting lines join the same cell line across methods); black bar = median across the four cell lines. Per-panel tags: Friedman test across methods (cell line as block; Benjamini-Hochberg FDR across the 12 properties) - no property differs significantly across methods. Variance decomposition (two-way ANOVA): cell line explains a median 98.8% of the variance per property versus 0.4% for fragmentation method.

**Supplementary Figure S16.**
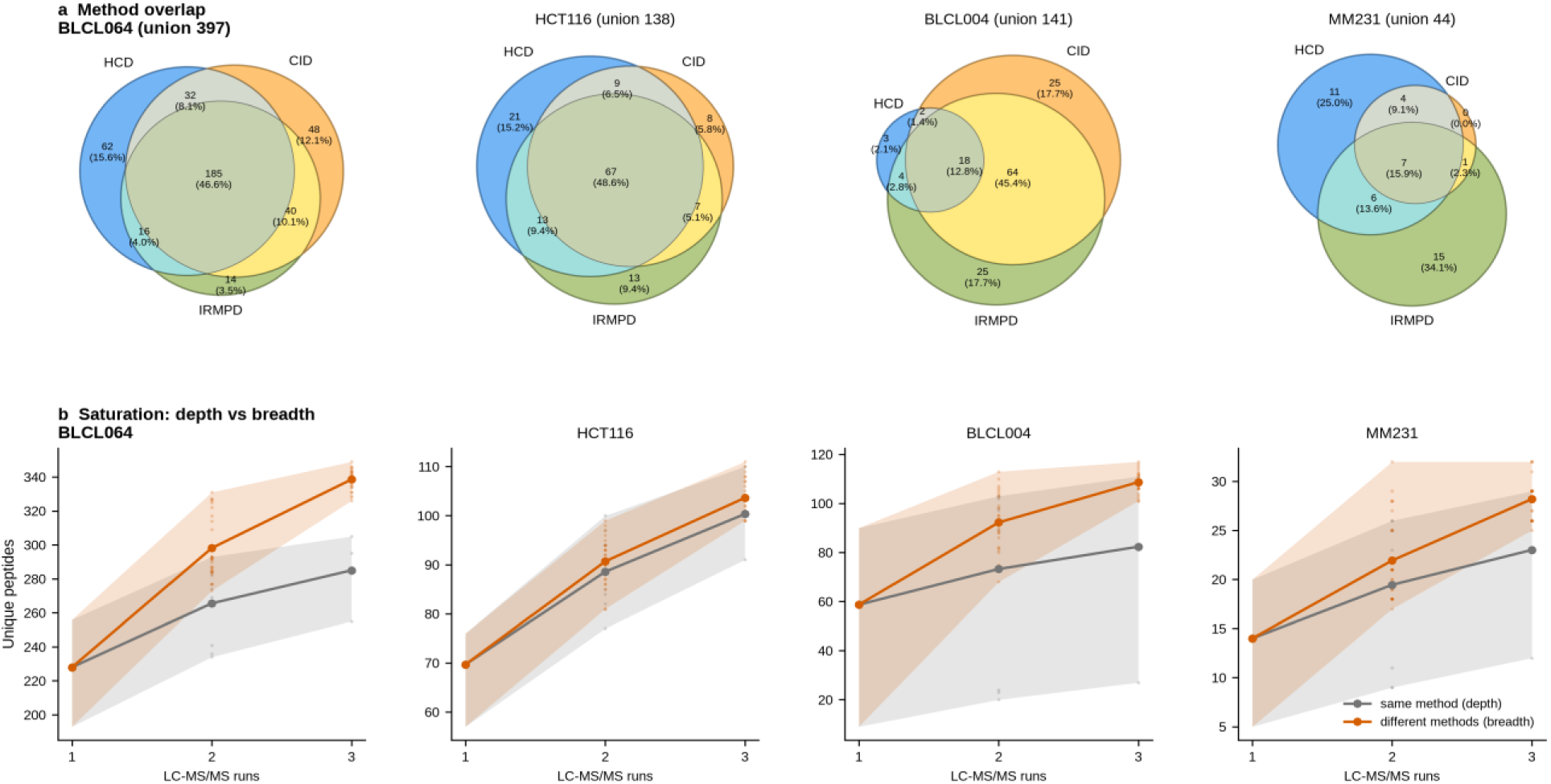
Non-redundant sampling and saturation analysis, cryptic database. Panels as in Figure 9. Cryptic unions are small (44-397 peptides per cell line) and dominated by the all -method shared core; the mean cross-method allocation exceeded the mean single-method triplicate in all four cell lines (+18.8% BLCL064, +32.0% BLCL004, +3. 3% HCT116, +22.5% MM 231).

**Supplementary Figure S17.**
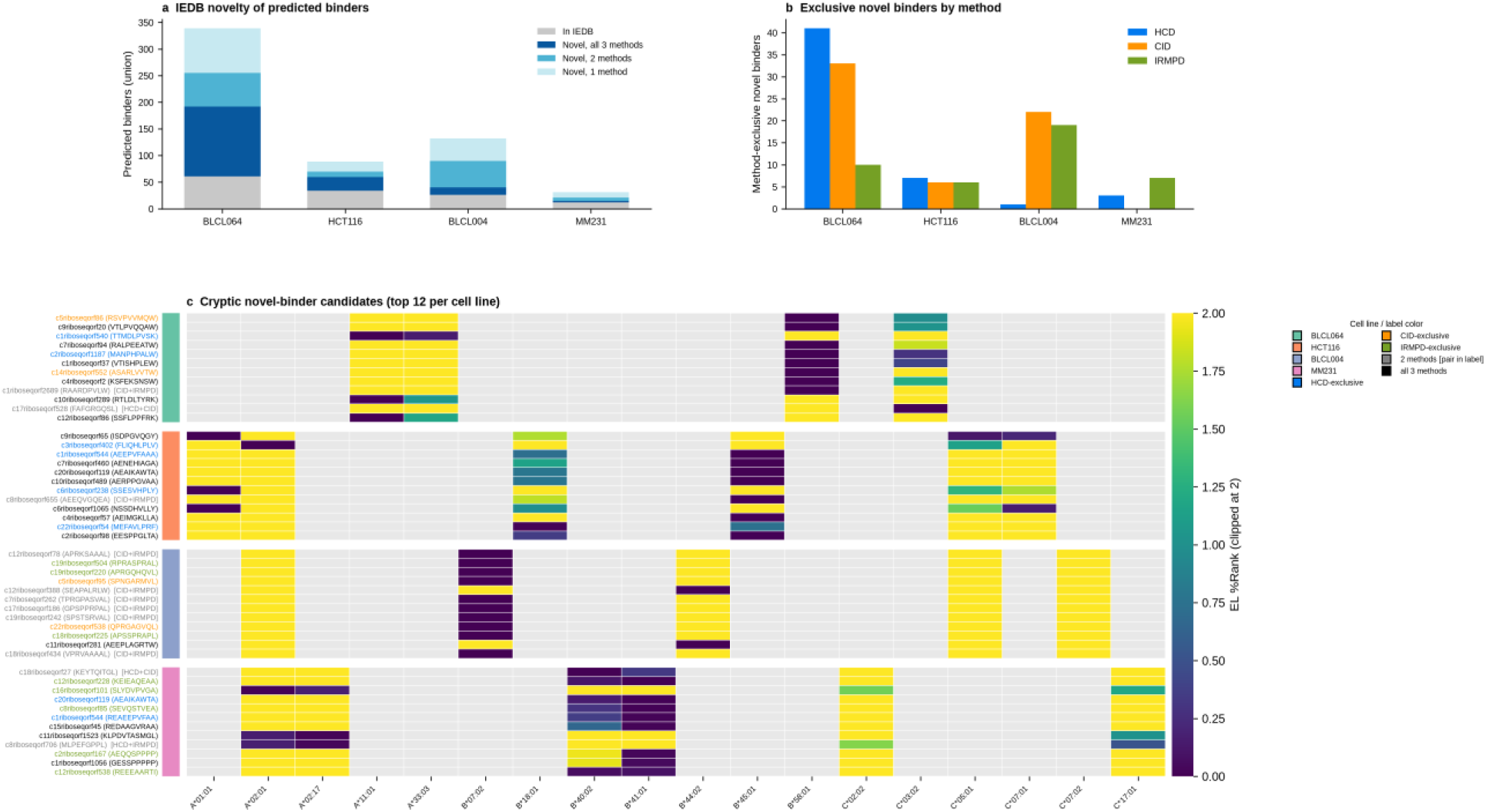
Novel cryptic HLA ligands (cryptic database). Panels as in Figure 10. 42.9-81.0% of cryptic peptides per cell line x method were absent from IEDB, yielding 458 novel predicted binders, 155 of them method-exclusive. No cryptic novel binder mapped to the curated cancer-antigen panel, so panel c shows the top 12 cryptic novel binders per cell line by best EL %Rank.

**Supplementary Figure S18.**
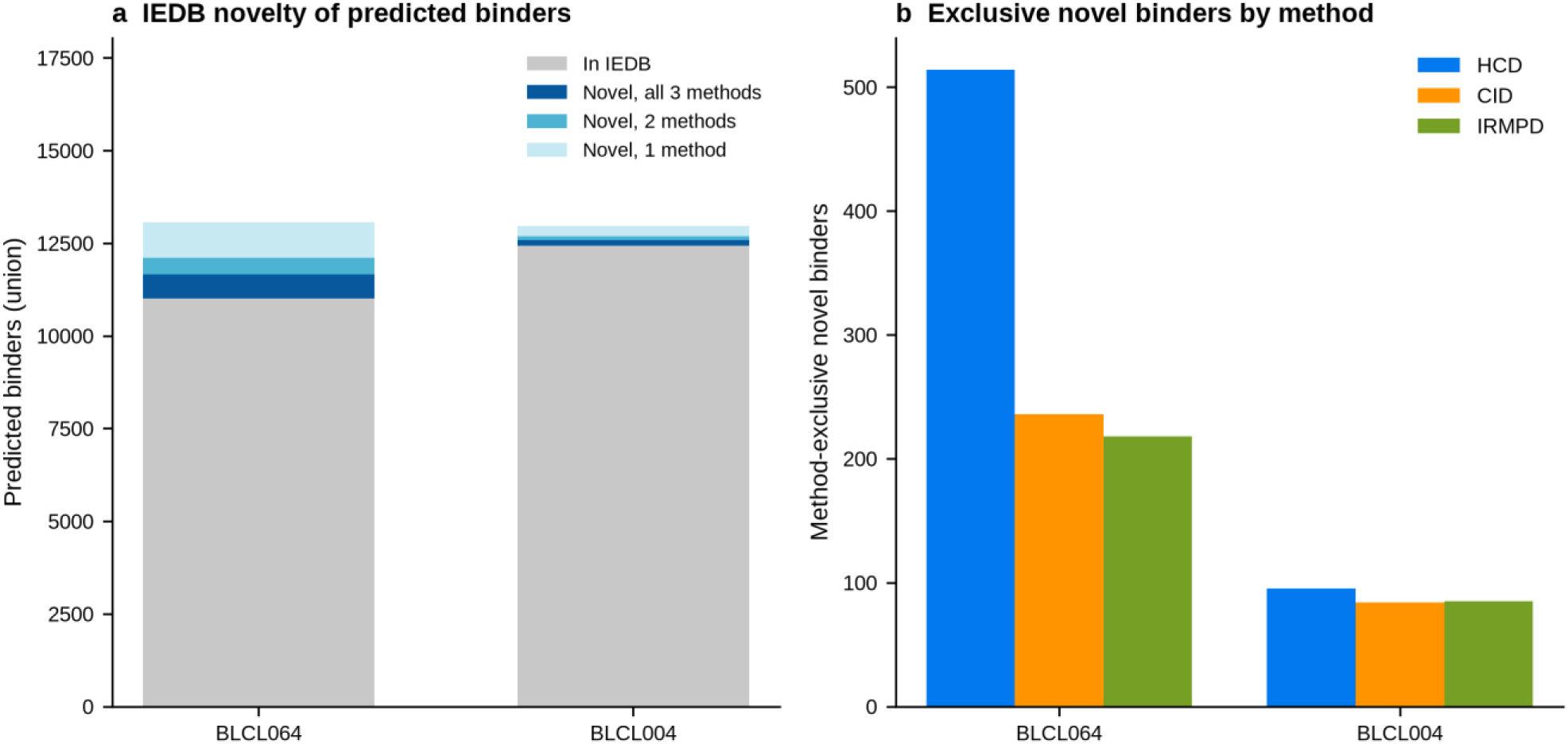
IEDB novelty and method-exclusive novel binders in the non-malignant reference lines. Panels as in Figure 10a,b, but for BLCL064 and BLCL004. (a) Predicted binders (union across methods) partitioned into in-IEDB (grey) and novel (blue shades, by number of detecting methods). (b) Method-exclusive novel binders per cell line, colored by detecting method.

**Supplementary Figure S19.**
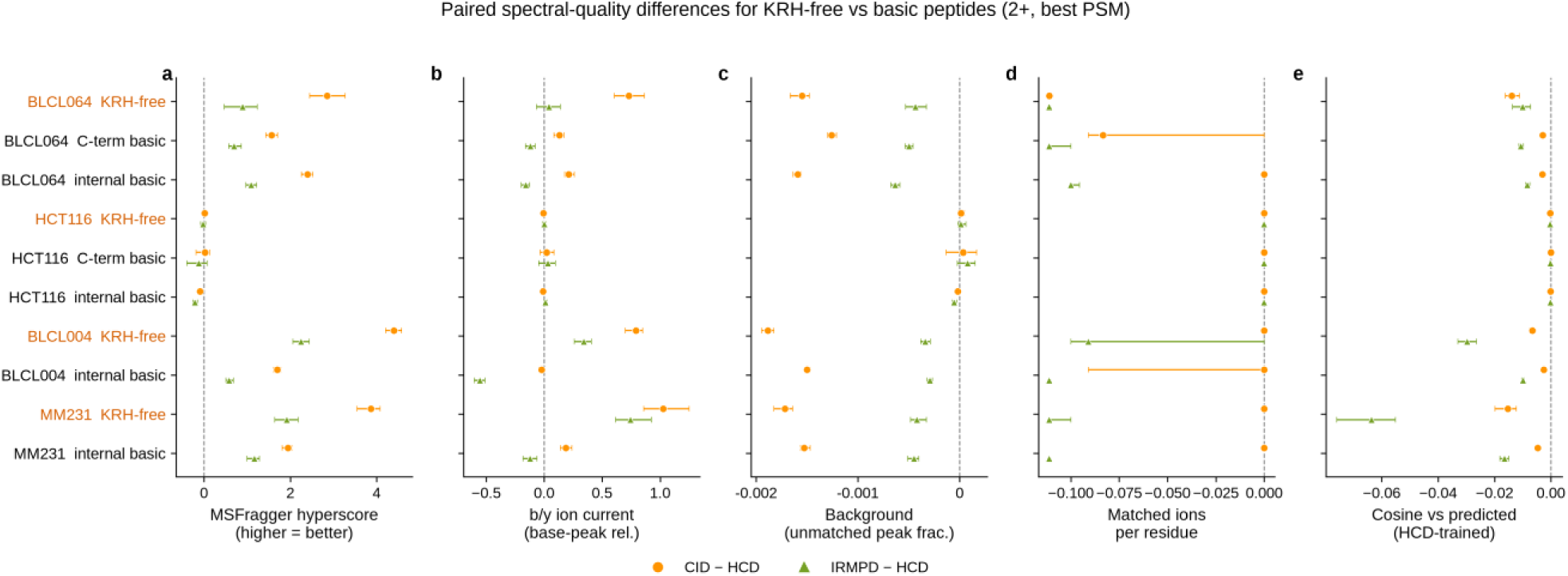
Paired spectral-quality differences for KRH-free versus basic peptides across all primary metrics. Each row is one cell l ine x residue-class combination (KRH-free rows in orange); class rows with fewer than 20 paired peptides are omitted (hence no C-terminal-basic rows for BLCL004 and MM231). Peptides identified by both compared methods were paired by sequence (best charge-2+ PSM per peptide per method); points are median paired deltas for CID - HCD (orange circles) and IRMPD - HCD (green triangles), error bars are 95% bootstrap confidence intervals, and the dashed line marks zero (no difference from HCD). Columns, left to right: (1) MSFragger hyperscore, the database-search score (higher = better); (2) summed b/y ion current, the sum of base-peak-relative intensities of all matched b/y fragment ions; (3) background fraction, the proportion of spectrum peaks not matched to any theoretical b/y ion (negative delta = cleaner CID/IRMPD spectra); (4) matched ions per residue, the number of distinct matched b/y ions divid ed by peptide length; (5) cosine similarity between the observed spectrum and the spectrum predicted by the HCD-trained AlphaPeptDeep model. Positive deltas favour CID or IRMPD over HCD for all metrics except background fraction, where negative deltas favour CID or IRMPD.

**Supplementary Figure S20.**
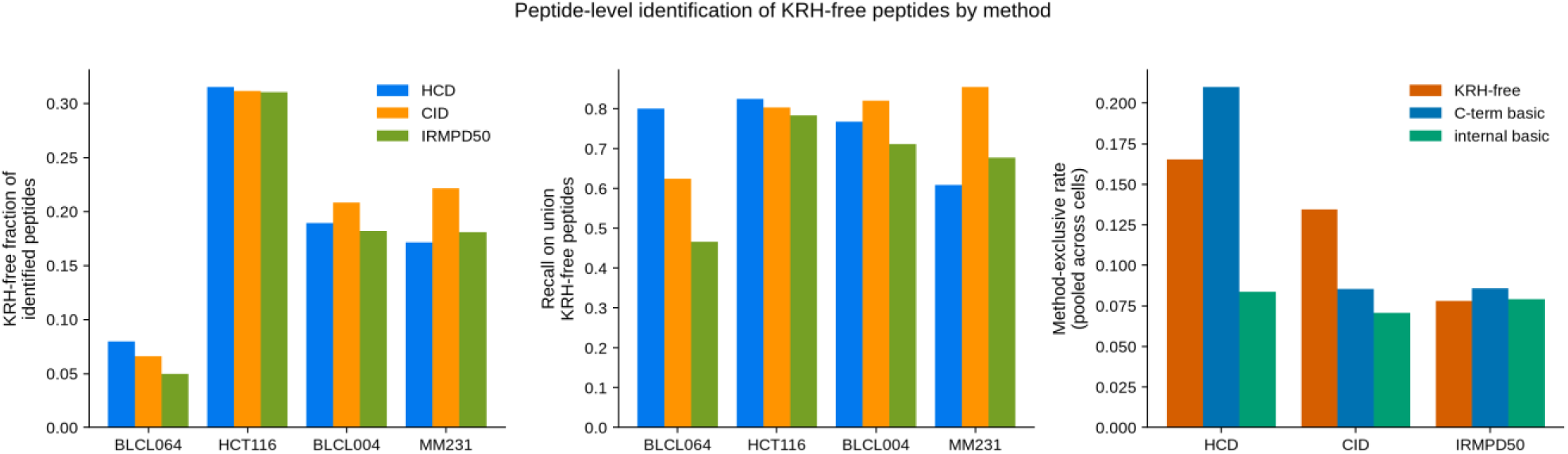
Peptide-level identification of KRH-free and basic-containing peptides per cell l ine and method. . Recall against the per-class union peptide set, and method-exclusive counts and rates, per residue class.

**Supplementary Figure S21.**
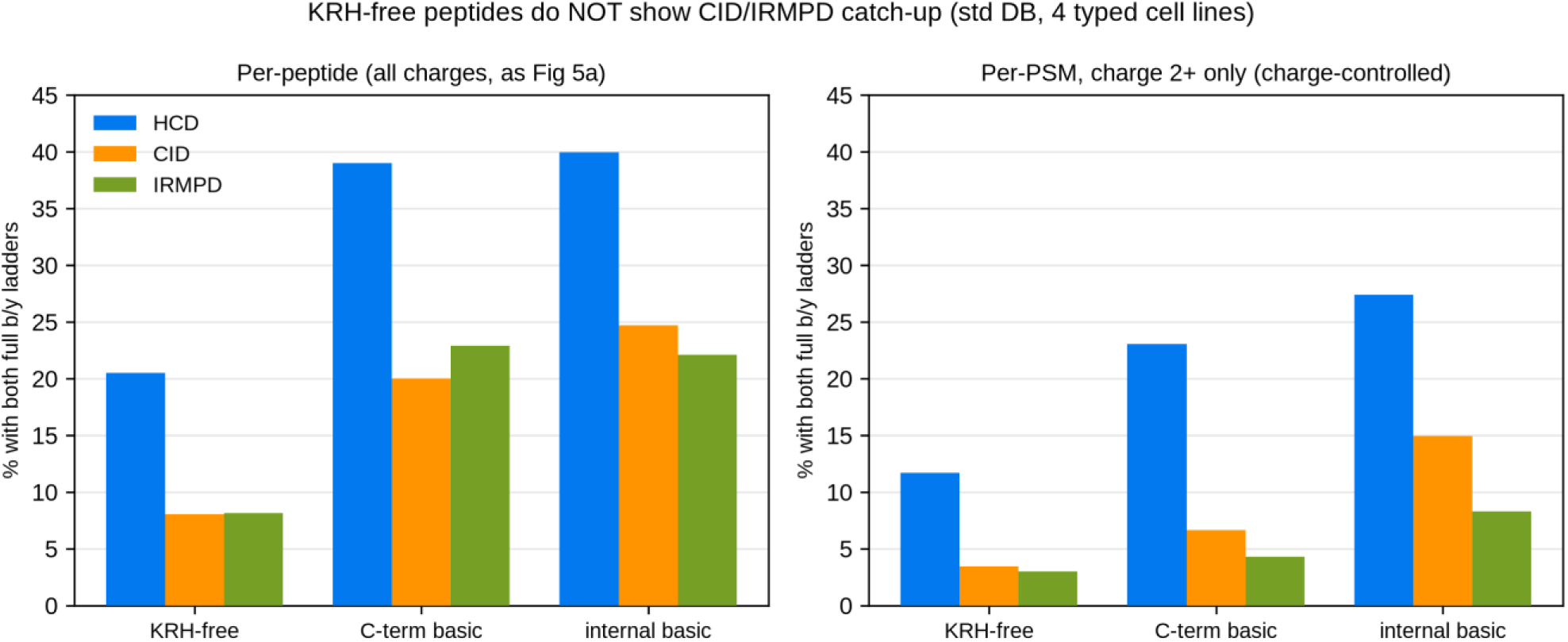
Fragment-ion ladder completeness by residue class and fragmentation method. Per-PSM completeness (fraction of theoretical b/y bonds observed) for KRH-free, internal-basic and C-terminal-basic peptides per cell line and method, including charge-state-matched (2+) comparisons. HCD retains the most complete ladders for all classes, including KRH-free peptides.

**Supplementary Table S 1. AlphaPeptDeep rescoring summary at 1% FDR (pooled triplicates per cell line for each method, both databases).** Rescued = peptides passing 1% FDR only after rescoring; lost = only before; gain (%) = 100 x (after - before) / before at the unique-peptide level.

**Supplementary Table S 2. Identification summary per cell line and method (standard database, pooled triplicates, 1 % FDR after AlphaPeptDeep rescoring).** Binder statistics cover 8-15mer peptides with NetMHCpan 4.2 predictions (EL %Rank < 2).

**Supplementary Table S 3. Ion-completeness matrix per cell line, method and database ( unique peptides).** Classes are mutually exclusive: both ladders = complete b - and y-ion ladders; full b/y only = one complete ladder with at least one gap in the other; gap-free only = every backbone bond covered by at least one matched ion without a complete ladder. The rem ainder to 100% has at least one uncovered bond. Median coverage = per-peptide median percentage of backbone bonds covered. All rows are restricted to 9mer predicted binders (best EL %Rank < 2), matching Figures 6 and S3.

**Supplementary Table S4.. Per-allele detection-share statistics by fragmentation method (standard database).** For each cell line x allele: size of the predicted-binder repertoire (union across methods), fraction detected by each method, and the CID - HCD and IRMPD- HCD share differences (percentage points) with two-proportion z-test p-values and Benjamini-Hochberg FDR per cell line.

**Supplementary Table S5.. Composition of common, rescued and lost peptide sets per method and database (median of per-cell-line values across the four HLA-typed cell lines; N peptides = pooled total).** Binder (%) = NetMHCpan 4.2 EL %Rank < 2 for the best allele, including supplementary predictions for peptides absent from the primary prediction tables.

**Supplementary Table S6.. Novel predicted binders (absent from IEDB, EL %Rank < 2 ) per cell line, broken down by detecting-method set (standard database).** Strong = EL %Rank < 0.5. Cancer-Ag = novel binders derived from the curated cancer-antigen panel. Full peptide-level lists are provided as Supplementary Tables S7 (standard) and S8 (cryptic).

**Supplementary Table S7.. Full list of novel predicted binders (absent from IEDB, EL %Rank < 2), standard database, including cancer-antigen annotations.**

**Supplementary Table S8.**. **Full list of novel predicted binders (absent from IEDB, EL %Rank < 2), cryptic database, including cancer-antigen annotations.**

**Supplementary Table S9.**. **Rescoring with foreign and pooled donor models versus the per-dataset own model (standard database; unique peptides, 1% FDR after rescoring).** Best foreign donor = single-donor model with the highest peptide yield; Master = model fine-tuned on all five cell lines including HeLa elastase. Master/own (%) = relative difference in unique peptides.

**Supplementary Table S10. KRH-free fraction of eluted ligands by HLA allele group and cell line. Allele groups: A2-supertype alleles, A3-supertype alleles, other; assignment by best MHCflurry EL rank.**

**Supplementary Table S 11. Paired spectral-quality comparison (CID-HCD and IRMPD-HCD) by residue class and cell line.** Per-peptide deltas of the best 2+ PSM per peptide; median delta with 95% bootstrap CI, rank-biserial correlation (rbc), Wilcoxon signed-rank p and Benjamini-Hochberg adjusted p (p_adj). Primary metrics flagged.

